# A short 3’UTR motif regulates gene expression in bilaterians

**DOI:** 10.1101/2023.11.15.567165

**Authors:** Ana Eufrásio, Joana Azevedo, Joana Machado, Alexandre Ferreira, Ana Moutinho, Filipe Henriques, Ana Jesus, Joana Tavares, Isabel Pereira-Castro, Joana Teixeira, Mafalda Araújo, Pedro B.P. Pinto, José Bessa, Alexandra Moreira

## Abstract

The mechanisms of gene expression regulation are essential for cell identity and function, and their disruption usually leads to human disease. The 3’ untranslated region (UTR) of mRNA contains important regulatory elements of gene expression, including upstream sequence elements (USEs) that are cis-regulatory sequences localized upstream of polyA signals (PAS). One of the best functionally characterised USEs is located in the 3’UTR of the Drosophila’s polo gene, which disruption leads to critical phenotypes in adult flies. In this work we found that the USE of the Drosophila’s polo gene (DplUSE) is also found in in the 3’UTR of vertebrate genes, including zebrafish, mouse and human genes, showing higher levels of conservation than the whole 3’UTR sequence. Using reporter assays, we show that DplUSE is able to increase gene expression in vitro in human cell lines and in vivo in zebrafish embryos. Importantly, in humans, the DplUSE containing genes are enriched for genes associated to serious diseases such as Congenital abnormalities and Malignant neoplasms, illustrating the potential of this sequence to modulate genes with relevant biological functions and related with human health. Concomitantly, when sequestering the molecular machinery that operates at the DplUSE using a dominant negative strategy, we show that this is enough to dysregulate DplUSE containing genes in human cells and disrupt proper embryo development in zebrafish. Aiming to understand the molecular mechanism operating at the DplUSE, we identified three RNA binding proteins (RBP) that specifically bind to the DplUSE in vertebrates. Importantly, one of such RBPs is PTBP1, the vertebrate orthologue of the fruit fly’s RBP Heph, that was demonstrated to be required for the DplUSE function in Drosophila. To test if PTBP1is essential for the DplUSE function, as observed in Drosophila, we depleted PTBP1 from human cells and observed a downregulation of the expression of DplUSE containing genes, demonstrating that the molecular mechanisms that operate at DplUSE are ultra-conserved. Finally, we explored if variants in DplUSE consensus could be associated to human disease. We found a reported single nucleotide polymorphism (SNP; rs3087967) that is associated with malignant tumor of colon and generates an ectopic consensus of DplUSE in the 3’ UTR of the tumorigenic POU2AF2/C11orf53 gene. We further show that this ectopic DplUSE motif causes a gain-of-function in vivo in zebrafish gut cells, suggesting its involvement in colon cancer development. These results show that a short motif present in the 3’UTR of genes from phylogenetically distant bilaterians, from fruit flies to humans, control genes’ expression through an ultra-conserved mechanism involving RBPs binding and its dysregulation might impact in human disease.

## Introduction

The 3’ untranslated region (3’UTR) of the mRNA is defined by cleavage and polyadenylation of the pre-mRNA and has a central function in regulation of gene expression. This level of regulation is achieved by the existence of specific sequences in the 3’UTR that bind RNA binding proteins (RBPs), or that are targeted by miRNAs, controlling the fate of the mRNA (reviewed in ^1–8)^. One of the most important regulatory mechanisms found in the 3’UTR are the polyadenylation signals (PAS), that are conserved among fish, flies, mammals, and worms^9^. Similarly to mammals, the most common PAS in zebrafish (Danio rerio) is the canonical AAUAAA, which is present in 62.3% of all genes, and also contains the GU/U-rich downstream sequence^10^. However, about half of pre-mRNAs do not contain a canonical PAS (AAUAAA)^11,12^ and yet, they are efficiently cleaved and polyadenylated. Usage of non-canonical PAS is enhanced and fine-tuned by the presence of auxiliary cis-elements in the pre-mRNA, such as the upstream sequence elements (USEs) that are localized upstream of the PAS. Therefore, USEs are fundamental for the proper control of gene activity. In humans, USEs have been characterized as sequences that activate PAS selection and 3’ end formation efficiency by acting as an extra-platform for the recruitment of trans-acting factors, such as RBPs, with an important function in diseases^6,13–26^. Other examples of USEs have been identified in other species such as the fruit fly (Drosophila melanogaster). In the 3’UTR of the Drosophila’s gene polo, which codes for a key cell cycle kinase^27^, two polyadenylation sites originate two distinct mRNAs^28^. Upstream of polo’s proximal PAS there is a highly conserved 28-nucleotides USE, which has a function in increasing Polo protein levels at the kinetochores^14^. Previous bioinformatic analysis has shown that the most conserved region of polo’s USE (TTGTTTTT; DplUSE) is present upstream of a PAS in 2.7% of all human genes^14^. Additionally, we have previously shown that human DplUSEs are more prevalent upstream of weaker PAS than the canonical AATAAA, including in the human genome, suggesting their role in enhancing PAS efficiency in human genes, as it occurs in fly genes. Reinforcing this hypothesis, a sequence similar to DplUSE, having one nucleotide mismatch (TTGT[G/T]TTT) has been found in the 3’UTR of the human gene prothrombin F2, being demonstrated its role as a regulatory sequence of gene activity and its potential link to human disease^29^. All these results suggest that the DplUSE might be present in vertebrate genomes and its genes’ regulatory function activity might be highly conserved, a question that is yet to be addressed.

Here we show that the DplUSE sequence in present in the 3’ UTR of several genes of zebrafish, mouse and humans, suggesting a conserved role of the DplUSE sequence in the regulation of genes activity. To further understand the DplUSE role, we have performed in vivo reporter assays in zebrafish using the DplUSE in the 3’UTR of the GFP reporter gene, showing that DplUSE increases the expression of GFP in several different tissues. Additionally, we demonstrated that microinjection of DplUSE RNA in one-cell stage zebrafish embryos leads to several abnormalities during embryonic development, possibly by sequestering the molecular machinery that operates in the DplUSE, as might be the case of specific RBPs. Indeed, when the DplUSE is over-expressed in human cells, genes that contain the DplUSE sequence are downregulated, further supporting the dominant negative effect of the ectopic expression of DplUSE. These results suggest that overall, the molecular machinery that operates in the DplUSE is required for the proper early embryonic development of a vertebrate species as the zebrafish. To better understand the molecular mechanism of the DplUSE, we have explored the binding of candidate RBPs to the DplUSE sequence, showing that HuR/ELAVL1^30–35^, hnRNPC^20,36^ and PTBP1^37–40^ RBPs bind to DplUSE. Importantly, we further show in human cells that when depleting PTBP1, the orthologue RBP that in the fruit fly was observed to be required for the function of the DplUSE, several genes that contain the DplUSE sequence are downregulated, suggesting that there is an ultra-conserved molecular mechanism controlling gene expression via the DplUSE sequence. Finally, we searched for SNPs that might affect the DplUSE sequence and are associated to human disease. We found that the rs3087967 SNP associated with malignant tumor of colon is located in the 3’UTR of the oncogene C11orf53/POU2AF2^41^ and creates an ectopic consensus of DplUSE, causing a gain of function of the mRNA and suggesting its involvement in colon cancer development. Collectively, we provide in vivo evidence for the function of DplUSE in zebrafish, its conserved mechanism of action in zebrafish and humans as a regulator of gene expression and the potential implications of the DplUSE consensus in human disease.

## Results

### A fruit fly upstream sequence element is conserved in the 3’ UTR of humans, mouse and zebrafish genes

In a previous study, we have identified a USE sequence (DplUSE) located in the 3’UTR of the cell cycle gene polo from the fruit fly (Drosophila melanogaster)^14^. We found that this sequence is essential to maintain Polo protein levels required for cell cycle progression^14^, suggesting that this USE could represent a regulatory mechanism for gene activity. To understand if the DplUSE sequence could be present in vertebrate species we have developed a bioinformatic script to identify all the genes that contain in their 3’UTR the DplUSE sequence (TTGTTTT) located at a maximum distance of 450 nucleotides (nt) upstream of the weak ATTAAA polyadenylation signal (Figure 1A).Our bioinformatic search identified the presence of the DplUSE in 2110 human genes, 1648 mouse genes and 714 zebrafish genes, with subgroups sharing orthologues between species (Figure 1B). From the 2110 human genes that contain DplUSE, we observed a significant enrichment for diseases such as congenital abnormalities, colorectal cancer and malignant neoplasms (MNeoplasms), comparing with the control genes (Figure1C). Importantly, 31 orthologue genes in human, mouse and zebrafish share the presence of the DplUSE in their 3’UTR (Figure 1B and D). From these 31 orthologue genes, we verified a significant enrichment for congenital abnormalities, MNeoplasms, and for nervous system diseases, comparing with the control genes (Figure S1A). Altogether, these results highlight the potential of the regulatory function of DplUSE in human diseases. Considering this, we then explored whether the DplUSE sequence could show a higher level of conservation than the respective 3’UTR region, when aligned with multiple vertebrate genomes. The results showed that indeed the DplUSE motif is significantly more conserved, when compared to the remaining 3’UTR, both in the 31 orthologue genes dataset, as in the 2110 human genes (Figure 1E and Figure S1B and C). Next, we evaluated the degree of conservation of each nucleotide of the DplUSE motif in vertebrates based on the multiple alignments of 100 vertebrate species and measurements of evolutionary conservation using phyloP^42^. For this analysis we extended 3 nucleotides (N) upstream and downstream of DplUSE. Using the DplUSE motifs identified in the 2110 human DplUSE containing genes, we observed that the TTT located in the 5th to 7th position of the DplUSE motif (TTGTTTTT) present highest values of conservation while the T located in the 4th position DplUSE motif (TTGTTTTT) presented the lowest value of conservation. Similar results were observed when analysing the DplUSE motif present in the 31 genes subset (Sup Fig1D). These results suggest that the DplUSE motif might accommodate some nucleotide variants. Nevertheless, using the multiple DplUSE human sequences aligned with the genome of key vertebrate species (mouse, alligator and zebrafish), we found that the most constant consensus was the TTGTTTTT motif, with the exception of the alignments with zebrafish, which motif was TTCTTTC (Sup.Fig.1F). Analyses of the gene ontology (GO) terms related to biological processes demonstrated that for Homo sapiens DplUSE-containing genes are involved in dynamic processes such as cellular migration, adhesion and communication, and mRNA processing (Figure S1 D). For Mus musculus, DplUSE-containing genes GO terms include regulation of epithelial to mesenchymal transition and cell response to glucose, and for Danio rerio, DplUSE-containing genes are involved in mitotic cell cycle process, protein ubiquitination and several developmental processes (Figure S1 E and F). We then asked if there were common GO terms that are highly enriched (F.E ≥ 1.5) between the three organisms (Figure S1G). Although there are no common GO terms for the three species all together, we found that 10 out of 22 human GO terms are common to mouse, 2 are common to zebrafish GO terms; and 4 out of 14 zebrafish GO terms are common to mouse (Figure S1 G and H). These GO terms include cell cycle related funstions such as “cell division” (human and mouse), “chromosome organization” (mouse and zebrafish), “mitotic cell cycle process” and “mitotic cell cycle” (human and zebrafish), which is consistent with the role of the fruit fly gene polo, from where the DplUSE has been originally identified, suggesting that DplUSE might have conserved regulatory functions in cell cycle control. GO terms for other important cell biology terms as “positive regulation of cell migration” and “mRNA processing” have been also identified.

**Figure 1.**
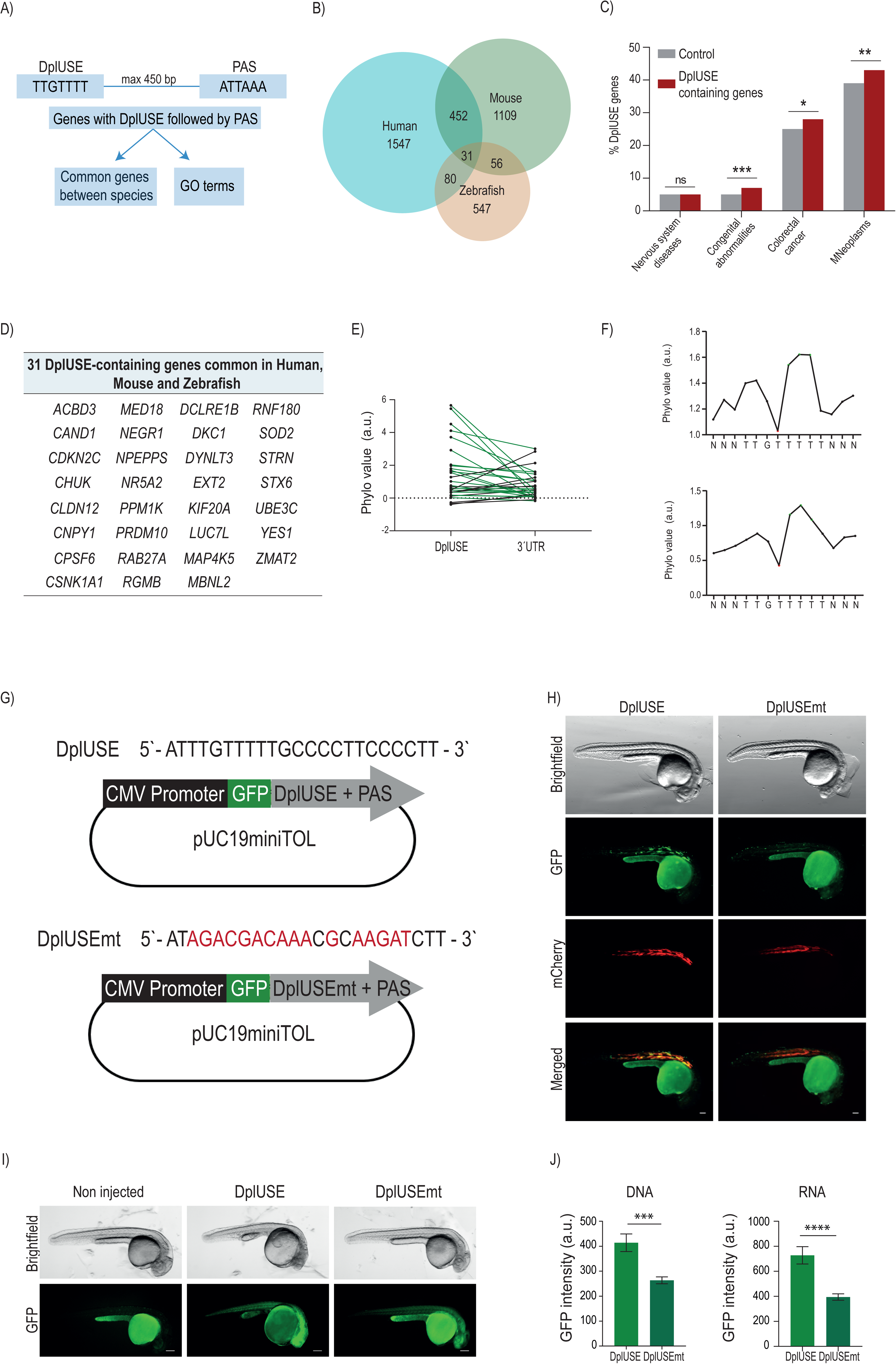
The DplUSE sequence is conserved across human, zebrafish and mouse genes and increases GFP expression in a post-transcriptional manner in zebrafish. (A) Workflow used to obtain the Gene Ontology (GO) terms related to biological processes of the common DplUSE-containing genes between the human and zebrafish transcriptomes obtained from an R-based bioinformatic script, which identified all the genes that contained, in the 3’ UTR, the most conserved region of the USE sequence (TTGTTTT) located upstream of the polyadenylation signal (ATTAAA), at a maximum separating distance of 450 base pairs (bp). (B) Venn Diagram representing: the number of DplUSE-containing genes found in human (2110), zebrafish (714) and mouse (1648). 80 USE-containing genes are common between human and zebrafish while 31 are common between the 3 species. (C) Representative graph showing the percentage of the 2110 DplUSE-containing human genes that are associated with the respective disease (DisGenet data source). (D) The 31 USE-containing genes that are common between the 3 species: human, mouse and zebrafish. (E) Representative graph of the average value of the extracted conservation values from each nucleotide within the USE and 3’UTR sequence, for each one of the 31 ortholog genes (data source:100vertebrates). (F) Representative graph showing the average conservation value for each nucleotide within the 2110 DplUSE containing human genes. (G) Schematic representation of pUC19miniTOL-GFP-USE and pUC19miniTOL-GFP-USEmt plasmids used to microinject one-cell stage zebrafish embryos. The black boxes correspond to the cytomegalovirus (CMV) promoter region; the green boxes correspond to the GFP reporter gene; and the grey boxes represent the USE sequence, or the USE sequence mutated in 16 nucleotides, followed by the polyadenylation signal (PAS)-ATTAAA. (H) Representative images from 24hpf zebrafish embryos microinjected with pUC19miniTOL-GFP-USE/pUC19miniTOL-GFP-USEmt with *Tol2* mRNA and mylz:mCherry vector. (I) Representative images from 24hpf zebrafish embryos microinjected with GFP-USE RNA and GFP-USEmt RNA. (J) Representative graph for the GFP expression quantification from microinjected embryos with pUC19miniTOL-GFP-USE and pUC19miniTOL-GFP-USEmt (left); Representative graph for the GFP expression quantification from microinjected embryos with *GFP-USE* and *GFP-USEmt* RNAs; Statistical significance was determined by χ2 test with Fisher correction or by two-tailed unpaired t-test. ***p<0.001; **p<0.01; *p<0.05; ns p>0.05. Images were acquired with Leica M205(scale bar = 100 μm) and the GFP expression was quantified for at least 25 embryos in each.

### The DplUSE sequence increases protein expression in a vertebrate model system

To understand if the DplUSE might act as a gene’s activity regulatory sequence in vertebrates, increasing gene’s activity as observed in the fruit fly, we have developed two expression vectors, pUC19miniTOL-GFP-DplUSE and pUC19miniTOL-GFP-DplUSEmt. These constructs are ubiquitous expression vectors containing the DplUSE sequence, or the DplUSE sequence mutated in 16 nucleotides (DplUSEmt), subcloned downstream of the GFP coding sequence and assembled in a Tol2 transposon (Figure 1F), suitable to perform transgenesis reporter assays in zebrafish embryos. Microinjection of these constructs in one-cell stage wild-type (WT) embryos, simultaneously with Tol2 mRNA and another transposon that drives expression of mCherry in the muscle (control of transgenesis), showed that, upon 24 hours post-fertilization (24 hpf), embryos microinjected with GFP-DplUSE DNA exhibited more intense GFP fluorescence, while embryos microinjected with GFP-DplUSEmt DNA demonstrated a more diffused and faded fluorescence (Figure 1G). Quantification of GFP intensity (Figure 1I, left) revealed that embryos microinjected with GFP-DplUSE DNA showed a significantly higher expression than when microinjected with GFP-DplUSEmt DNA. These results demonstrate that the DplUSE is able to enhance GFP expression in a vertebrate species as in the fruit fly, demonstrating its conserved function from arthropods to vertebrates.

To better understand if this activating effect takes place at a transcriptional or post-transcriptional level, GFP-DplUSE and GFP-DplUSEmt RNA were in vitro transcribed and microinjected in one-cell stage zebrafish embryos. Despite the fact that the fluorescence in these embryos was far more diffuse than the one observed for embryos microinjected with the expression vectors (Figure 1H), it is still possible to notice and quantify a higher GFP expression in embryos microinjected with GFP-DplUSE RNA than with GFP-DplUSEmt RNA (Figure 1H and 1I, right), in a similar manner to the results obtained upon the microinjection of GFP-DplUSE expression vector (Figure 1G and 1I, left). These results indicate that in zebrafish the DplUSE sequence increases gene expression in a post-transcriptional manner, which is compatible with the mechanistic action of the DplUSE sequence found in the fruit fly, where it was observed to be dependent on the binding of RBPs^14^.

### DplUSE RNA recruits protein factors essential for the correct expression of genes involved in zebrafish development and in human cells function

To better characterize the function of the DplUSE sequence in controlling gene’s activity, a transgenic GFP-DplUSE zebrafish line was generated using the Tol2 transposon system^43^. Although the expression of GFP is driven by a ubiquitous promoter (CMV), the stable transgenic line exhibits GFP expression mainly in the notochord, muscle, pituitary gland and eye (Figure 2A and 2B), suggesting a possible role of the DplUSE in regulation of gene expression in a tissue and cell-type specific manner. Specifically, confocal imaging demonstrates the clear and intense pattern of GFP expression in the notochord and muscle (Figure 2B), suggesting that the DplUSE sequence might have tissue specific regulatory functions. To address the function of the DplUSE and the respective molecular machinery that operates in this sequence, we considered a dominant negative approach, induced by ectopic introduction of DplUSE RNA. In this case we postulate that factors such as RBPs might be sequestered by the ectopic DplUSE RNA, affecting therefore the expression of DplUSE controlled genes. To test this hypothesis, we have microinjected DplUSE RNA in transgenic GFP-DplUSE embryos and we quantified the GFP expression, in comparison with the microinjection of DplUSEmt RNA (USE sequence mutated in 17 nt) (Figure 2C). We found that the microinjection of USE RNA was able to decrease GFP expression, contrasting with the microinjection of USEmt RNA at 5 dpf, particularly in the notochord and in the muscle (Figure 2D to F), therefore corroborating the dominant negative effect hypothesis. These results were consistent at 48 hpf, where it was also observed a decrease in GFP expression in the notochord (Figure S2A). Further observation of the imaging revealed that the microinjection of DplUSE RNA also affected the consistency of the GFP expression, leading to the development of patchy GFP expression, a phenomena that occurred less frequently upon microinjection of DplUSEmt RNA at 5 dpf (Figure 2D and G) and at 48 hpf (Figure S2A and S2C).

**Figure 2.**
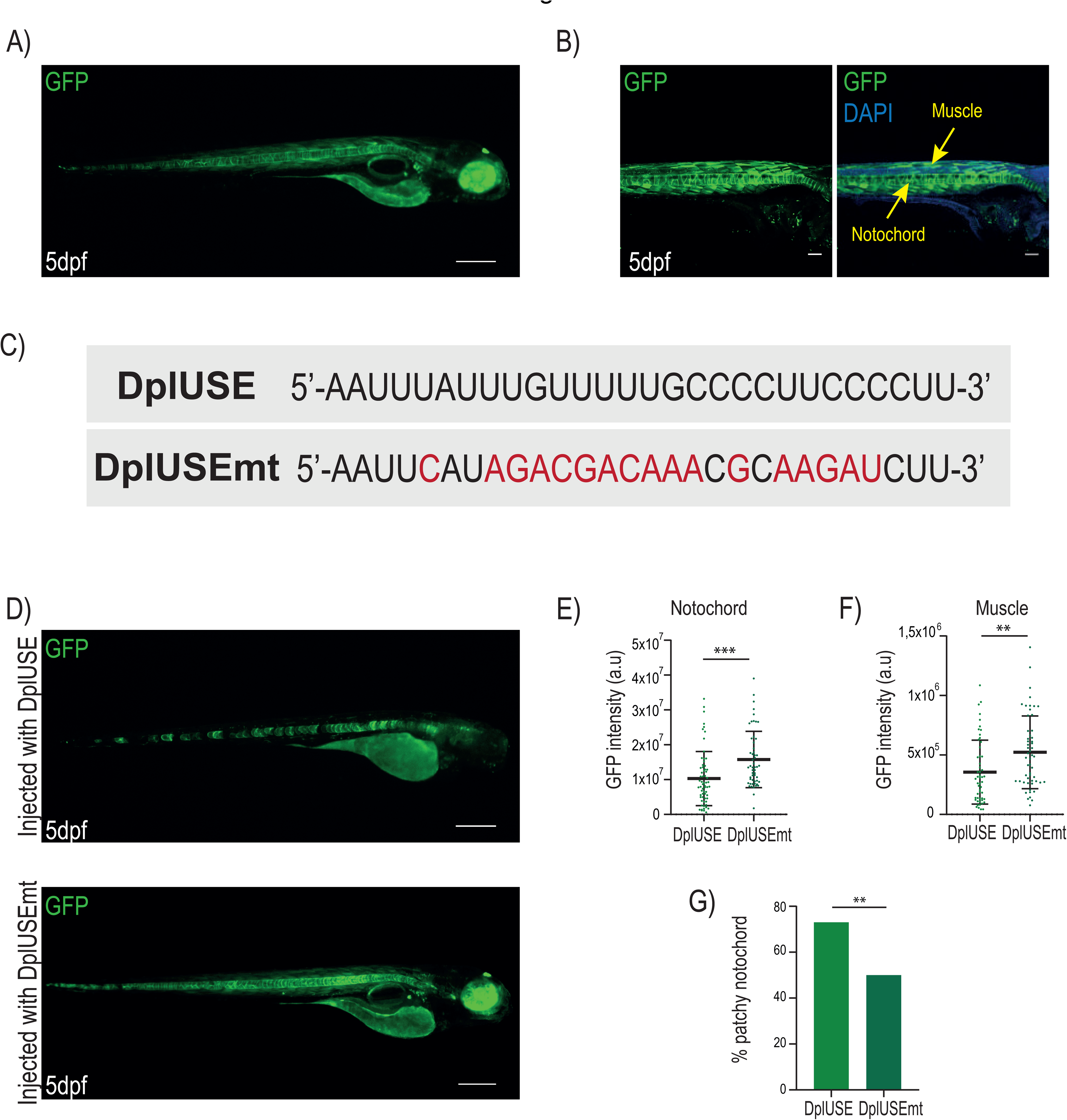
Microinjection of Dpl*USE* RNA causes a decrease in GFP expression in transgenic GFP-USE animals. (A) Representative image of a 5dpf GFP-DplUSE transgenic line. (B) Confocal images of transgenic GFP-USE zebrafish, stained with the nuclear marker DAPI, showing GFP expression in the notochord and muscle (Scale bar= 10 μm). (C) Dpl*USE* RNA and Dpl*USE*mt RNA sequences used to microinject transgenic GFP-DplUSE zebrafish embryos. (D) Representative image of GFP expression in transgenic *GFP-DplUSE* zebrafish embryos microinjected with Dpl*USE* RNA or USEmt RNA, at 5 dpf. Images were acquired using Leica M205. (E) Quantification of GFP expression in notochord (n=64 Dpl*USE*; n=54 Dpl*USEmt*) and (F) in muscle (n=53 Dpl*USE*; n=54 Dpl*USEmt*). (G) Quantification of the percentage of transgenic *GFP-*Dpl*USE* embryos that showed a patchy expression of GFP in the notochord (73%,n=69 Dpl*USE*;50%,n=.59 Dpl*USEmt*) Statistical significance was determined by χ2 test with Fisher correction or by two-tailed unpaired t-test. ***p<0.001; **p<0.01; *p<0.05; ns p>0.05. (Scale bar = 100 μm).

Next, we have injected DplUSE RNA and focused on developed phenotypes. At 5 dpf, zebrafish larvae microinjected with DplUSE RNA exhibited microphthalmia, a condition where either one or both eyes are abnormally small or absent^44^. When the eye size was quantified, a significant decrease was observed, when compared to larvae microinjected with DplUSEmt RNA (Figure 3A and B). This phenotype was also observed at 48 hpf (Figure S2D and S2E). Apart from these eye phenotypes, a careful evaluation of 24 hpf larvae allowed us to identify other important phenotypes affecting the notochord, heart, head, late gastrulation and tail, that are more prevalent in DplUSE RNA than with DplUSEmt RNA injected embryos (Figure 3C-E). At 5 dpf, malformations in the notochord, head, tail and swim bladder where also detected (Figure S3A, B and C). To better understand the mechanisms that might be causing these phenotypes, we inquired if apoptosis was increased by performing immunohistochemistry staining with anti-cleaved caspase-3 polyclonal antibody (pAb) and analysed the presence of cleaved caspase-3 fluorescence signal in 48hpf embryos microinjected with DplGFP-USE and DplGFP-USEmt RNAs. We observed a more prevalent presence of cleaved caspase-3 staining in embryos microinjected with DplGFP-USE RNA, when comparing with not injected embryos and embryos injected with DplGFP-USEmt RNA (Figure 3F and G), suggesting that, in part, these phenotypes are caused by apoptosis. Overall, these results show that ectopic DplUSE RNA might work as a dominant negative, interfering with the normal DplUSE regulatory functions, as show using the GFP-DplUSE reporter line. In addition, ectopic DplUSE RNA causes several developmental abnormalities, possibly by sequestering RBPs that bind to the DplUSE mRNA of endogenous USE-containing genes, interfering with the expression of these genes and others that might also depend on the function of these RBPs and which expression is necessary for correct embryo development.

**Figure 3.**
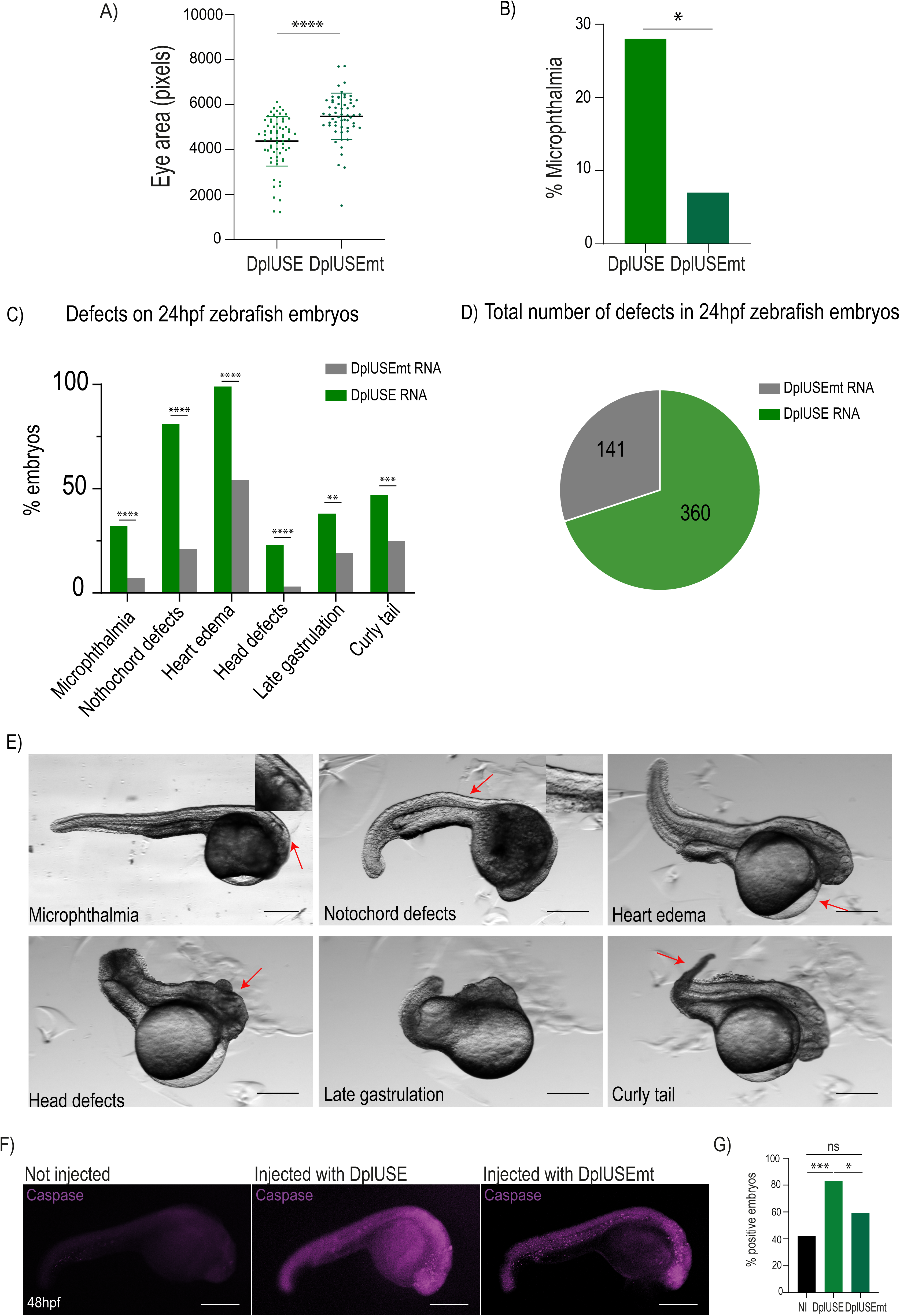
Microinjection of Dpl*USE* RNA in one-cell stage zebrafish embryos induces the development of several defects. (A) Representative graph of the quantification of eye size (pixels) in microinjected embryos with Dpl*USE* and Dpl*USEmt* RNAs, at 5dpf. (n=69 Dpl*USE*; n=58 Dpl*USEmt*) (B) Representative graph of que quantification of the percentage of microphthamia in microinjected embryos with Dpl*USE* and Dpl*USEmt* RNAs, at 5dpf (28%, n=68 DplUSE; 7%,n=58 Dpl*USEmt.* (C) Quantification of the several defects developed 24h upon the microinjection of *GFP-DplUSE* RNA (in green) and *GFP-*Dpl*USEmt* RNA (in grey) in one-cell stage zebrafish embryos. (D) Pie chart representing the total number of defects developed upon the microinjection of Dpl*GFP-USE* RNA and *GFP-*Dpl*USEmt* RNA. (E) Representative images showing the various abnormalities upon microinjection of *GFP-* Dpl*USE* RNA at 24 hpf, indicated with red arrows. (F) Representative images of not injected embryos and in embryos injected with Dpl*USE* and Dpl*USE* RNAs, at 48hpf, showing in purple, the cleaved anti-caspase-3 staining. (G) Representative graph showing the percentage of embryos with consistent appearance of apoptotic cells, characterized by the presence of anti-caspase-3 staining (42%,n=39 Not injected; 83%,n=33 Dpl*USE*;59%, n=33 Dpl*USEmt*). Statistical significance was determined by χ^2^ test with Fisher correction or by two-tailed unpaired t-test. ***p<0.001; **p<0.01; *p<0.05; ns p>0.05 Images acquired with Leica M205.Scale bar = 100 μm.

To understand if the DplUSE sequence shares similar regulatory mechanisms in human cells as observed in the fruit fly and zebrafish, we started by asking whether the DplUSE would have an activating effect in gfp expression in HeLa cells, using reporter constructs. Transfection with the pUC19miniTOL-GFP-DplUSE and pUC19miniTOL-GFP-DplUSEmt constructs show that DplUSE is also able to enhance the expression of GFP, showing a ∼2-fold increase in GFP expression with the USE-containing construct in comparison to the USEmt construct (Figure 4A). These results recapitulate the ones observed in zebrafish and suggest that the DplUSE gene regulatory mechanisms are conserved in vertebrates. Furthermore, similar to what we observed in zebrafish (Figure 2D), we reasoned that the DplUSE transcribed from the transfected plasmid may also act as a dominant negative, titrating the molecular machinery that operates at the DplUSE, resulting in a decrease of expression of genes that are under the control of DplUSE. Thus, after transfection, the mRNA levels of nine DplUSE-containing genes (KIF20A, EXT2, CLDN12, MBNL2, MED18, NR5A2, CSNK1A1, DKC1 and MAP4K5), selected from the 31 genes identified by the bioinformatic script that are common to human, mouse and zebrafish (listed in Figure 1D) were quantified by RT-qPCR (Figure 4B-J). Our show that the transfection of GFP-DplUSE leads to a significant decrease in the expression of most of these genes in comparison with GFP-DplUSEmt (Figure 4B-J). Thus, our results show that the ectopic expression of DplUSE might act as a dominant negative, downregulating the expression of genes that contain the DlpDplUSE and that are associated with cancer, such as KIF20A which is a tumour-associated antigen involved in the glioma cell growth and cell survival, Claudin12, which is overexpressed in colorectal carcinomas, NR5A2, which negatively regulates cell proliferation and tumour growth in nervous system malignancies, CSNK1A, which is a negative regulator of canonical Wnt signalling pathway and of oncogenic RAS-induced autophagy and Cdkn2c, which suppresses tumorigenesis, further supporting the conserved role of the DplUSE sequence as regulator of gene expression in vertebrates.

**Figure 4.**
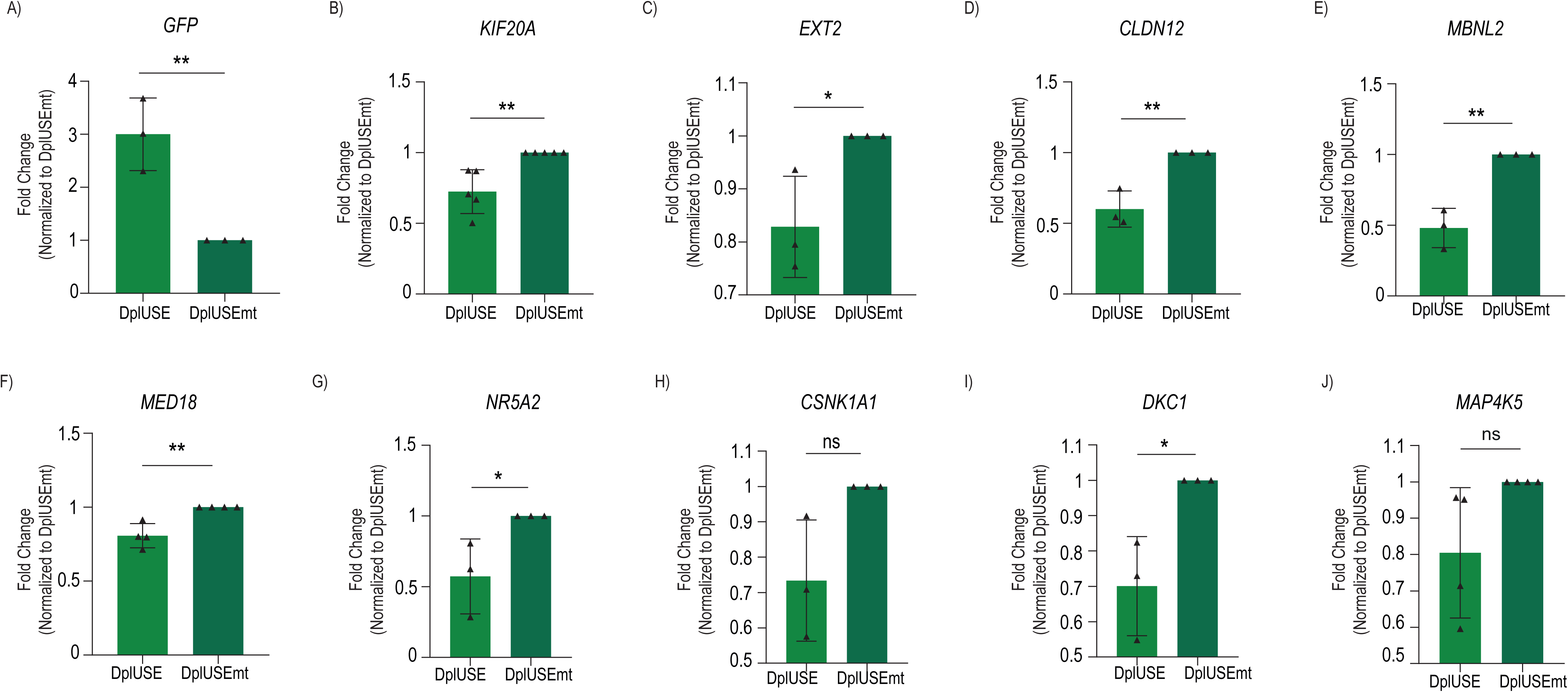
The DplUSE increases *GFP* expression and modulates DplUSE-containing genes in human cells. (A) RT-qPCR showing that the mRNA levels of *GFP* increase by 3-fold upon transfection with the pUC19miniTOL-GFP-DplUSE plasmid (DplUSE) when comparing with the transfection with the pUC19miniTOL-GFP-DplUSEmt plasmid (USE mut). mRNA levels of *KIF20A* (B), *EXT2* (C), *CLDN12* (D), *MBNL2* (E), *MED18* (F), *NR5A2* (G), *CSNK1A1* (H), *DKC1* (I) and *MAP4K5* (J) were assessed by RT-qPCR. Apart from *CSNK1A1* and *MAP4K5*, transfection with the pUC19miniTOL-GFP-USE plasmid decreases the mRNA levels of the USE-containing human genes. The data is presented as the mean ± SD of at least three independent experiments and analysed by two-tailed unpaired Student’s t-test. ***p<0.001; **p<0.01; *p<0.05; ns p>0.05.

### A bilaterian ultraconserved molecular mechanism operates in the DplUSE RNA

Next, we aimed to better understand the molecular mechanism of the DplUSE sequence in controlling gene’s activity. We reasoned that, because of its nature as a USE located at the 3’UTR of genes, DplUSE could be binding RBPs, that have been shown to be players of USEs function^22,26,40,45,46^To test this hypothesis, and to determine if human proteins bind to the DplUSE, constructs containing the DplUSE or DplUSEmt were used to perform in vitro transcription generating radioactively labelled pre-mRNAs that were incubated with human HeLa cells nuclear extracts to perform UV cross-linking assays. The results show that multiple proteins assemble onto the DplUSE pre-mRNA (Figure 5B, lane 1) and that mutation of the USE sequence leads to a decrease in the binding (Figure 5B, lane 2). To determine the binding specificity of these proteins to the DplUSE RNA motif, UV cross-linking competition assays were performed, where increasing amounts of unlabelled DplUSE or DplUSEmt RNA were added to the reaction mix (Figure 5C). The results showed that DplUSE RNA unlabelled competitor (1 pmol, 50 pmol and 150 pmol-Figure 5C, lane 2, 3 and 4, respectively) significantly inhibits the assembly of the ∼35 kDa, ∼40 kDa and ∼60 kDa proteins onto the radioactive labelled DplUSE pre-mRNA. This competition is DplUSE sequence specific, as addition of the same amounts of unlabelled DplUSEmt RNA (1 pmol, 50 pmol and 150 pmol – Figure 5C, lane 5, 6 and 7, respectively) did not cause any noticeable changes in the proteins bound to DplUSE pre-mRNA. These results also reinforce the dominant negative mechanism observed when overexpressing the DplUSE in zebrafish embryos (Fig. 2, 3 and 4) and human cells (Fig. 4).

**Figure 5.**
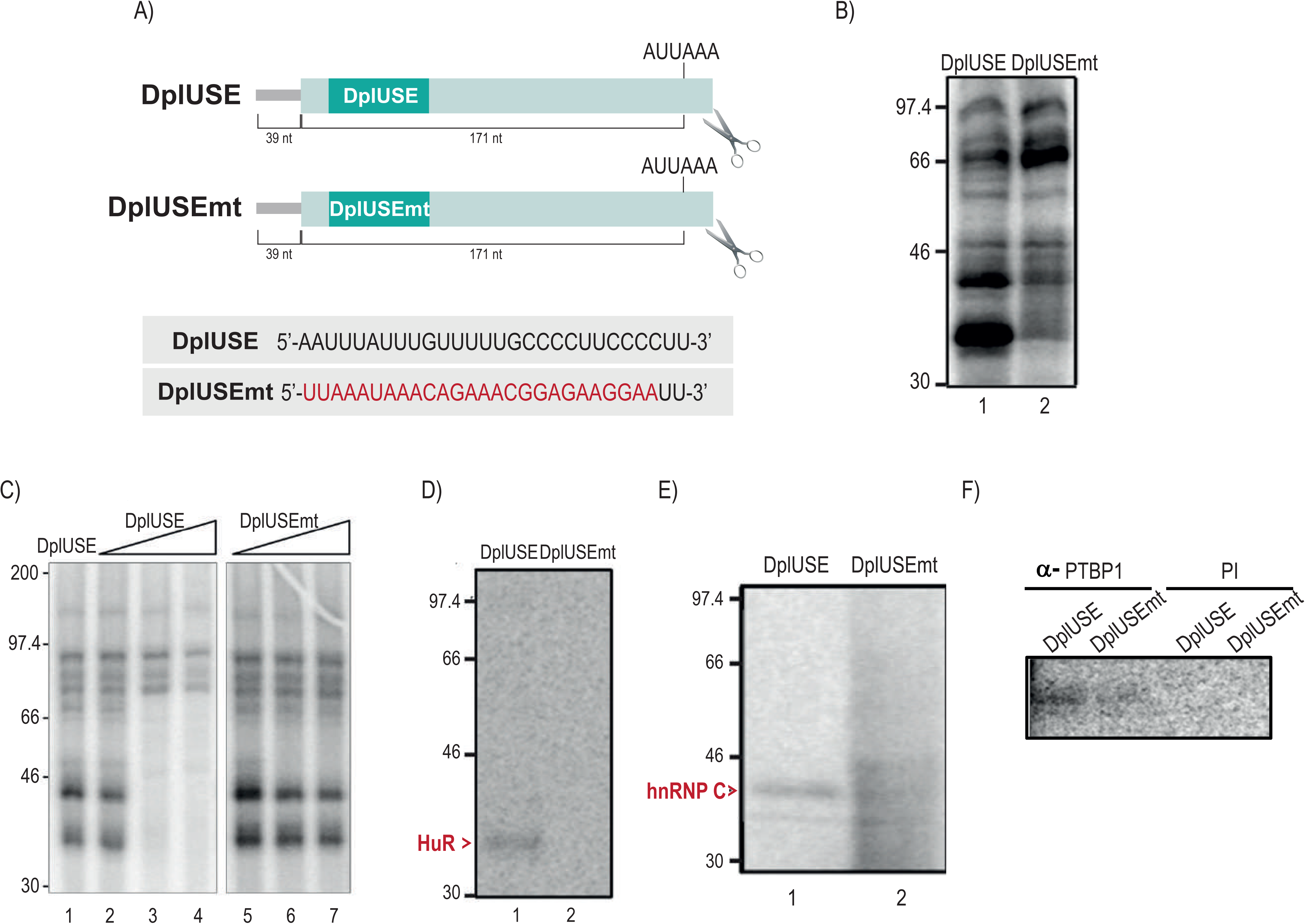
HuR, hnRNP C and PTBP1 bind to the DplUSE sequence in a specific manner in human cells. (A) Schematic representation of the pre-cleaved pre-mRNAs used to perform the UV cross-linking assays. The DplUSE sequence and the DplUSE sequence mutated in 26 nucleotides (DplUSEmt) is presented. The cleaved pre-mRNAs were obtained by linearizing the plasmids with restriction enzymes in proximity to the cleavage site (portrayed by the pair of scissors). (B) UV cross-linking assay using the pre-cleaved pre-mRNAs DplUSE and DplUSEmt incubated with HeLa cells nuclear extracts. Asterisks define the bands correspondent to the proteins that undergo a decrease in the binding upon the mutation of the DplUSE sequence to DplUSEmt. (C) UV cross-linking competition assay using pre-cleaved pre-mRNAs DplUSE and DplUSEmt with HeLa cell nuclear extracts (lane 1) and an increasing amount of unlabeled DplUSE (lane 2-4) and DplUSEmt (lane 5-7) pre-mRNAs. One (lane 2, 5), 50 (lane 3,6) and 150 pmols (lane 4,7) of unlabeled oligonucleotides were used to compete with the labeled pre-mRNAs. (D-F) Immunoprecipitations assays performed subsequently to UV cross-linking of pre-cleaved pre-mRNAs DplUSE and DplUSEmt with HeLa cell nuclear extracts. Immunoprecipitations were performed using HuR, hnRNP C and PTBP1 antibodies.

In the fruit fly, the RBP Heph was previously shown to bind to DplUSE RNA of the fly’s gene polo^14^ being required for the DplUSE regulatory functions. For this reason, we tested if the human orthologue of Heph, the RBP PTBP1, could bind to the DplUSE RNA in human cells, similarly to the fruit fly, supporting an ultra-conserved molecular mechanism operating at the DplUSE. To test this hypothesis, we performed immunoprecipitation assays after UV cross-linking. Results show that similarly to Heph, the human orthologue PTBP1 binds specifically to the DplUSE and not to the DplUSEmt control RNA (Fig. 5F). PTBP1 is a 57 kDa protein with multiple functions^47^ that acts as a regulator of alternative splicing^48,49^ with neuronal functions^39,50,51^ and as a polyadenylation regulator in virus, humans and Drosophila^14,20,37,40,45^.

To identify other RBPs that are binding to the DplUSE we analysed with detail the molecular weights of the proteins assembled onto the DplUSE pre-mRNA (Figure 6B, lane 1 and Figure 6C). We hypothesized that HuR/ELAVL1 and hnRNPC could be binding to DplUSE. HuR is a 32 kDa RBP, ubiquitously expressed in all human tissues that recognizes and binds to U-rich and AU-rich elements present in the 3’UTR of target mRNAs modulating their stability^30,31,34,52^. Furthermore, its fruit fly orthologue ElavL1 has been shown to promote 3’UTR extensions observed in the nervous system^35^. In addition, HnRNP C1/C2 isoforms (41 and 43 kDa, respectively) bind to polyU tracts and assemble on nascent transcripts and modulate several mRNA functions, including mRNA stability^36,53,54^. Using antibodies against these proteins it was possible to immunoprecipitate these RBPs after UV cross-linking using the DplUSE pre-mRNA and HeLa cell nuclear extracts (Figure 5D-F). In contrast, when the UV cross-linking assays were performed with USEmt pre-mRNA, the proteins were not as efficiently immunoprecipitated (Figure 5D). Overall, these results show that PTBP1, HuR and hnRNPC bind specifically to the DplUSE RNA in human cells, likely making part of the molecular regulatory mechanism operating at the DplUSE sequence.

**Figure 6.**
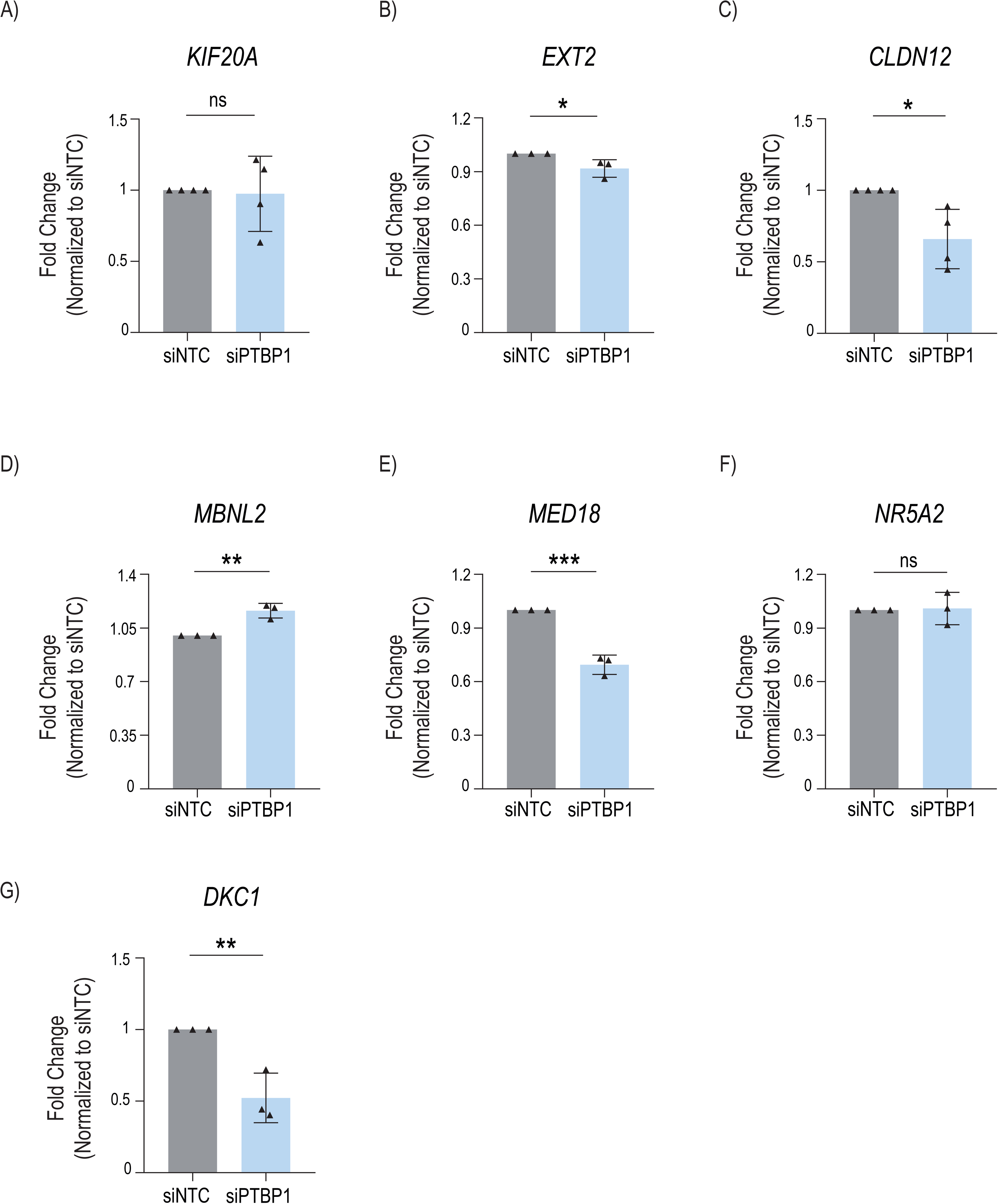
PTBP1 knockdown affects the expression of human DplUSE-containing genes. (A) PTBP1 knockdown was performed using siPTBP1 and DplUSE-containing genes mRNA levels were assessed by RT-qPCR. A siNTC pool was also transfected to be used as a control. *EXT2* (B), *CLDN12* (C), *MED18* (E) and *DKC1* (H) mRNA levels decreased upon *PTBP1* knockdown, like expected, but *MBNL2* (D) mRNA levels increased. For *KIF20A* (A) and *NR5A2* (F) the difference was not statistically significant. The data is presented as the mean ± SD of at least three independent experiments analysed by two-tailed unpaired Student’s t-test, *p < 0.05, **p < 0.01, ***p < 0.001 and ns – not significant.

To further demonstrate that the fruit fly’s polo sequence TTGTTTT is responsible for the ultra-conserved binding of RBPs to the pre-mRNA in human cells, we tested an expanded version of the fruits fly’s DplUSE sequence (AATTTATTTGTTTTTGCCCCTTCCCCTT). We also included specific mutations at the 5’ half of the sequence (USE-5’mt; AATT(C)AT(A)(G)(A)(C)(G)(A)(C)(A)GCCCCTTCCCCTT), at the 3’ half (USE-3’mt; AATTTATTTGTTTTT(A)(A)C(G)C(A)(A)(G)(A)(T)CTT) and both 5’ and 3’ together (USEmt-2; AATT(C)AT(A)(G)(A)(C)(G)(A)(C)(A)(A)(A)C(G)C(A)(A)(G)(A)(T)CTT; Figure S4A) and performed in vitro transcription and UV cross-linking assays using nuclear extracts from human Hela cells. These results show that proteins bind strongly only to the non-mutated extended DplUSE sequence and to the USE-3’mt sequence, therefore suggesting that the ultra-conserved bilaterian binding of RBPs is located at the DplUSE sequence (TTGTTTT; Figure S4A). To further validate these results, we performed similar assays splitting the expanded DplUSE sequence in three different fragments: DplUSE -(a), DplUSE -(b) and DplUSE -(c) (Figure S4A). Results show that the USE-(a) fragment and the expanded DplUSE sequence, the only tested fragments that contain the DplUSE sequence (TTGTTTT), bind strongly to proteins. Reinforcing the role of HuR and hnRNPC RBPs in the regulatory function of the DplUSE sequence, in both assays it was observed that the proteins which binding depends of the DplUSE sequence (TTGTTTT) have MWs similar to HuR and hnRNPC (Figure S4A).

### PTBP1 is part of the molecular mechanism modulating the expression of USE-containing genes

To investigate if PTBP1 is required for the control of the expression of DplUSE-containing genes, we performed PTBP1 depletion using siRNAs in HeLa cells. A knockdown of approximately 75% was obtained (Figure S5) and the expression of nine USE-containing genes (KIF20A, EXT2, CLDN12, MBNL2, MED18, NR5A2, DKC1) was quantified by RT-qPCR. These genes were selected by their potential to respond to the dominant negative effect of the ectopic expression of DplUSE (Figure 5). We observed that EXT2, CLDN12, MED18 and DKC1 mRNA levels decreased significantly upon PTBP1 knockdown, while KIF20A and NR5A2 were not clearly affected and MBNL2 mRNA levels increased (Figure 6). Our results indicate that PTBP1, as its fruit flýs orthologue Heph, modulate the expression of DplUSE-containing genes, supporting the existence of a regulatory mechanism that operates at DplUSE and that is conserved in bilaterians.

### A single nucleotide polymorphism in the DplUSE sequence is associated with increased cancer risk

In agreement with the mechanism of action of previously described USEs and the results of this study, the DplUSE RNA sequence acts as a platform for the recruitment of RBPs to regulate gene expression. Thus, modifications in the USE sequences may be involved in the deregulation of gene expression in the context of human disease. To investigate the potential impact of the DplUSE sequence in human diseases, we searched for the occurrence of single nucleotide polymorphisms (SNPs) overlapping with the DplUSE sequence (TTGTTTT) (Figure 7A). We found two genes containing a SNP associated with human diseases in the DplUSE sequence. One of these SNPs has been associated to the increased risk of development of Malignant tumor of colon (MTC) (rs3087967; POU2AF2/C11orf53^55^) and the other is associated with alanine aminotransferase blood levels after remission induction therapy (rs149940960; GREB1^56^) (Figure 7B). Of note, one of these genes, POU2AF2/C11orf53 has been recently described to act as a coactivator of POU2F3 to maintain chromatin accessibility, therefore having the potential to control genetic programs^41^. Additionally, it has been also shown that depletion of POU2AF2/C11orf53 using an in vivo xenograft model of small cell lung cancer, repressed tumour growth and delayed progression of disease in mice^41^, showing the tumorigenic potential of these gene in specific cellular contexts. Importantly, the MTC risk-associated variant (rs3087967) reflects the acquisition of an ectopic DplUSE consensus, compared with the respective non-risk variant. Therefore, in agreement with our results that show increased gene expression in the presence of DplUSE consensus in the 3’UTR of genes, we asked if the rs3087967 variant, in the context of the 3’UTR of the tumorigenic POU2AF2/C11orf53 gene, has the potential to increase its expression levels. To test this, we cloned the 3‘UTR of POU2AF2/C11orf53, containing the two variants (risk and non-risk) downstream of GFP, under the control of a CMV promoter. Using the Tol2 system, we performed mosaic transgenesis assays in zebrafish and analysed the larvae at 4 dpf. Quantifying the GFP levels in gut cells, we observed that the risk variant showed higher levels of GFP expression, when comparing with the non-risk variant (Figure 7C and D). These results suggest that the acquisition of an ectopic USE in the 3‘UTR of the tumorigenic POU2AF2/C11orf53 might lead to an ectopic expression of this gene and consequently, contribute to the development of MTC. These results indicate that the DplUSE may have important unidentified roles in the clinical context.

**Figure 7.**
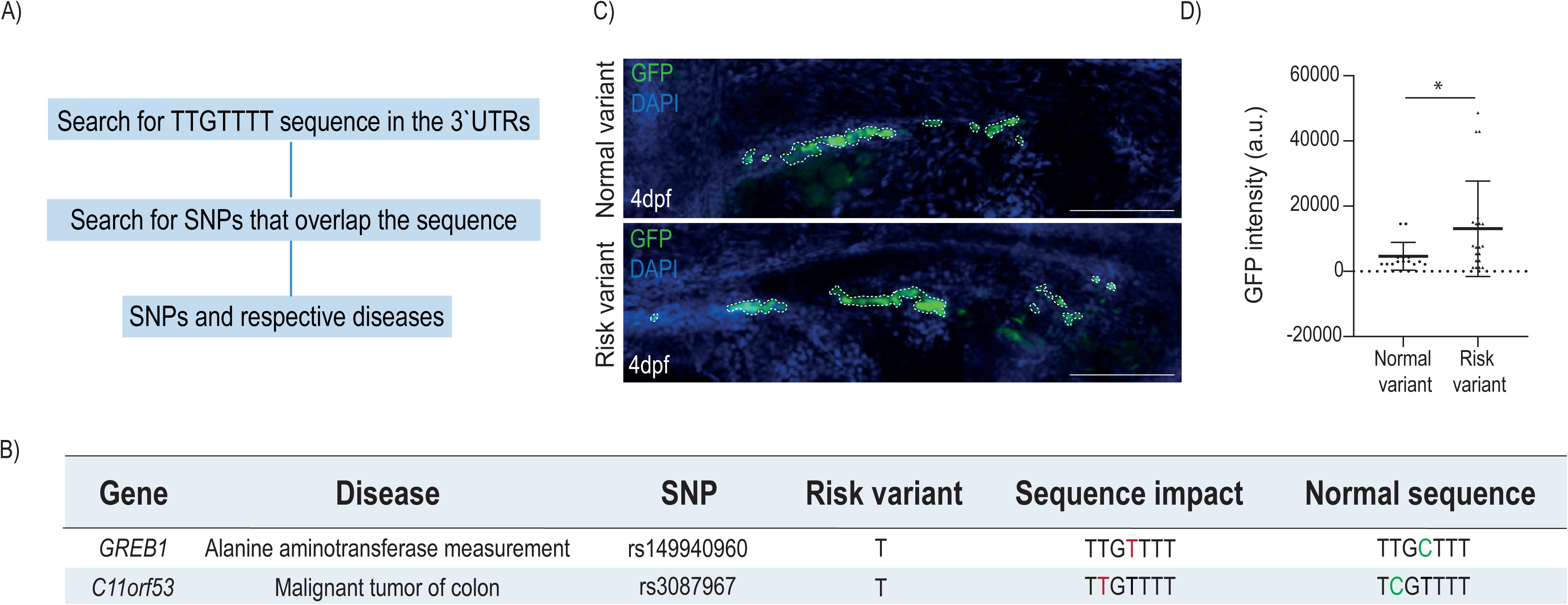
A SNP in *C11orf53* generates an ectopic consensus of the DplUSE which is associated with malignant tumour of colon. (A) Workflow used to identify SNPs that overlap with USE sequence and that had an associated disease. (B) DplUSE related SNPs with the respective gene, disease, sequence impact and normal sequence. (C) Representative confocal images of microinjected embryos with DplGFP-USE of C11orf53 with and without the presence of the risk variant, showing GFP expression in gut cells of 48hpf zebrafish embryos. The dashed lines represent different GFP expression domain within an embryo(D) Representative graph of the quantification of the GFP expression in the defined GFP expression domains (n=14, normal variant; n=20 risk variant). The images were acquired using the confocal microscope LeicaSP5II and the GFP expression quantified using IMARIS software. The statistical analysis was done by two-tailed unpaired Student’s t-test, *p < 0.05, **p < 0.01, ***p < 0.001.

## Materials and Methods

### Bioinformatic analysis

The NCBI RefSeq transcripts and the respective genes of Danio rerio (assembly danRer10), Mus musculus (assembly mm10) and Homo sapiens (assembly hg19) genomes were retrieved from UCSC browser [56]. The 3’UTR regions were selected and queried for the presence of the most conserved USE sequence (TTGTTTT) followed by pA1 (ATTAAA) spaced at a maximum of 450 bp, using an R-based script. Subsequently, the zebrafish genes were converted to human orthologs using BioMart^57^.

The Venn diagrams were made by using BioVenn. The Gene Ontology terms from human, zebrafish and mouse USE-containing gene lists were obtained using the GO Enrichment Analysis using Panther tools^58^. Only GO terms related to biological processes with a false discovery rate (FDR) <0.05 were considered. The terms were ordered by the fold enrichment (FE) and then by the FDR. The USE sequence data was intercepted by a dataset of SNPs described as linked to human diseases, available at the DisGeNET database^59^. Single-nucleotide polymorphisms (SNPs) that occurred in the USE sequence and that had a clinical significance were identified.

### Disease-association diseases and DplUSE vs 3’UTR sequence conservation analysis

After obtaining the coordinates for the most conserved USE sequence, we, subsequently extracted associated disease information from DisGeNet datasets. The disease data was sourced from human-curated information, combining data from UniProt, ClinVar, Orphanet, the GWAS Catalog, and CTD, using the Python programming language.

The 3’UTR coordinates regions were obtained from USCS genome browser and the phyloP values for each nucleotide from 100 vertebrate dataset track. This dataset represents multiple alignments of 100 vertebrates species and the measurement of evolutionary conservation using phyloP values These phyloP values serve as a metric for measuring evolutionary conservation at individual alignment sites. Positive scores indicate higher conservation, while negative values suggest lower conservation, relative to the evolution predicted under neutral drift.

### Oligonucleotides and Antibodies

The list of oligonucleotides and antibodies used in this work is in Supplementary Table 1.

### Plasmid constructs

pUC19mini-TOL-USEmt constructs contain the USE/USEmt sequences (from Drosophila melanogaster), USE normal variant/USE risk variant (from C11orf53) and the PAS (ATTAAA) located between both Tol2 arms^43^. *GFP-USE* and *GFP-USEmt* pre-mRNAs were synthesized by *in vitro* transcription using pCS2-GFP-USE and pCS2-GFP-USEmt constructs, respectively. pCS2-GFP-USE/pCS2-GFP-USEmt contains a CMV promoter located upstream of the GPF coding sequence and USE/USEmt sequence. In order to microinject GFP-*USE* and GFP-USEmt RNAs in the transgenic GFP-USE zebrafish embryos, *in vitro* transcription using pCS2-USE construct was performed. This construct contains the SP6 promoter upstream of the USE sequence.

pSPT19-*polo* wt, pGEM7-*polo* wt, pGEM7-*polo* wt-DSE, pSPT19-*polo* mt, pGEM7-*polo* mt or the pGEM7-USE1mt-DSE contain the USE/USEmt sequence located downstream of T7 or SP6 promoters, and they were constructed using the pBS-polo and pxb7 plasmids previously described^14^, and pGEM-7Zf(+) and pSPT19 as vector backbones.

### *In vitro* transcription

For synthesis of capped mRNA to microinject in zebrafish embryos, the template plasmids were initially linearized with appropriate restriction enzymes. In vitro transcription was performed incubating 1x transcription buffer, 5 mM DTT, 1 mM ATP, 1 mM UTP, 1 mM CTP, 0.5 mM GTP, 2.5 mM G(5’)ppp(5’)G RNA Cap Analogue (New England Biolabs), 4 μg DNA, 100 U Ribonuclease Inhibitor (NZYtech) and 60U of SP6 RNA polymerase (Thermo Fisher) for 2 h at 37°C. The DNA was digested by adding 10 U DNase I recombinant (Sigma) and incubating for one additional hour. Transcripts were purified by performing phenol/chloroform purification. For UV cross-linking assays, the USE pre-mRNAs were synthesized by performing *in vitro* transcription using pSPT19-*polo* wt, the pGEM7-*polo* wt or the pGEM7-*polo* wt-DSE constructs as templates. The USEmt pre-mRNAs were synthesized by *in vitro* transcription using the pSPT19-*polo* mt, pGEM7-*polo* mt or the pGEM7-USE1mt-DSE. The USE-(a), USE-(b) and USE-(c) pre-mRNAs were synthesized by *in vitro* transcription using constructs with pGEM-7Zf(+) as the parental vector. *In vitro* transcription was performed as previously described^40^, with minor modifications, by incubating 1 μg of linearized plasmids with 1x transcription buffer (Roche Applied Science), 10 mM DTT, 28.79 U of RNAguardTM Ribonuclease Inhibitor, 0.1 mM GTP, 0.5 mM CTP, 0.05 mM ATP and UTP, 1 mM Cap Analogue (Ambion), 8 μCi of α-^32^P-[ATP] and α-^32^P-[UTP] and 20 U of T7 RNA Polymerase or SP6 RNA Polymerase (Roche Applied Sicences). The reaction was incubated for 1 h at 37°C. The DNA was digested by adding 5 U of DNase I and incubating at 37°C for 15 min. The transcripts were purified using Illustra ProbeQuantTM G-50 Micro Columns according with the manufacturer’s instructions.

### UV cross-linking assays

These assays were performed as previously described^40^ with minor modifications. Briefly, a mixture containing 2 mM EDTA, 1 mM ATP, 20 mM CP, 11.5 U RNAguard^TM^ Ribonuclease Inhibitor, 2.5% (w/v) PVA and 40 μg/mL of tRNA was incubated with 100 cps of radiolabeled RNA and 5 μL of HeLa cell nuclear extracts. Reactions were incubated for 10 min at 30°C then 1 μL of 2.5 μg/mL tRNA was added. For competition assays 1, 50 and 150 pmoles of unlabeled competitor RNAs were added with labelled RNA. The reaction mixtures were irradiated twice with 96×10^4^ μJ/cm^3^ of UV light for 3 min, using the Hoefer UVC 500 Ultraviolet Crosslinker (GE Healthcare Life Sciences). RNase A was added to the samples, for 30 min at 37°C, to degrade unprotected RNA. Samples were boiled at 95°C for 5 min in a 2x SDS gel-loading buffer, in order to denature proteins, and separated by electrophoresis in a 10% SDS-PAGE. The gel was fixed a 10% (v/v) acetic acid, 10% (v/v) glycerol solution for 30 min at room temperature and then dried at 80°C under vacuum for 2 hours. The radiolabeled proteins bands were visualized by autoradiography.

### Immunoprecipitations

For immunoprecipitation of UV crossed-linked proteins with monoclonal antibodies 4F4 anti-hnRNP C (mouse) (kind gift from Gideon Dreyfuss, Howard Hughes Medical Institute, University of Pennsylvania School of Medicine) [61] and HuR (19F12, Thermo) 400 μL of 10% (v/v) protein A Sepharose CL-4B beads (GE Healthcare Life Sciences) in IP-2 buffer [50 Mm Tris pH 7.9, 50 mM NaCl, 0.1% (v/v) NP-40] was incubated with 40 μg of rabbit anti-mouse antibody (DakoCytomation) in a vertical wheel for 90 min at 4°C. The beads were washed 3 times with ice-cold IP-2 buffer. One UV cross-linking reaction and 2 μL of anti-hnRNP C or HuR was added to the beads and the mixture was rotated for 1 h at 4°C. The beads were subsequently washed 3 times and dried. For PTBP1 immunoprecipitation, a rabbit anti-PTBP1^60^ serum was used (generous gift from Chris W. J. Smith, Department of Biochemistry, University of Cambridge). To 100 μl of 50% (v/v) protein A sepharose-PBS, 20 μl of anti-PTBP1 and 600 μl of PBS were added, and the mix incubated for 1 hour at 4° C with gentle mixing. The beads were then washed with PBS and an UV crosslinking reaction was added and incubated overnight at 4° C. After incubation, the beads were washed two times with 800 μl of binding buffer I (20 mM Hepes pH 7.9, 150 mM NaCl, 0.05% (v/v) triton X-100) and then two times with binding buffer II (20 mM Hepes pH 7.9, 150 mM NaCl, 1% (v/v) triton X-100). As a non-specific control, an immunoprecipitation was performed using a rabbit pre-immune serum. 20 μL 2x SDS gel-loading buffer was added to the beads and the proteins were denatured at 95°C for 5 min and separated by gel electrophoresis in a 10% SDS-PAGE. The gel was fixed and dried at 80°C under vacuum for 2 h and the radiolabeled protein bands were visualized by autoradiography.

### Zebrafish husbandry and embryo culture

Adult wild-type Tuebingen (TU) zebrafish (*Danio rerio*) were maintained in a recirculating system under conditions approved by the i3S Animal Welfare and Ethics Committee and the Portuguese National Authority for Animal Health (DGAV). Fertilized eggs were kept at 28 °C in E3 medium supplemented with 0.001% 1-phenyl-2-thiourea to prevent pigmentation development^61^.

### Microinjection procedures, Zebrafish GFP-USE transgenic line establishment, GFP screening and quantification

One-cell stage AB WT embryos were microinjected with 3nL containing 25ng/ul *Tol2* transposase mRNA and 25ng/ul phenol/chloroform-purified reporter vectors: pUC19-miniTOL-GFP-USE, pUC19-miniTOL-GFP-USEmt and mylz:mCherry, being the latter used as internal control of transgenesis, driving mCherry expression specifically in muscle fibers of the zebrafish. Embryos were screened at 24hpf for GFP expression, using a fluorescence stereomicroscope. The images were acquired using Leica M205 and analyzed using ImageJ software^62^. For the GFP-RNAs microinjection, one-cell stage AB WT embryos were microinjected with 3nL containing 300ng/ul GFP-USE and GFP-USEmt RNAs. Embryos were screened and quantified at 24hpf for GFP expression as described before. Generation of the GFP-USE zebrafish transgenic line was performed using the *Tol2* transposon system. One-cell stage AB wild-type embryos were microinjected with 3nL containing 25ng/ul *Tol2* transposase mRNA and 25ng/ul phenol/chloroform-purified reporter vector (pUC19-miniTOL-GFP-USE). Embryos were screened at 48hpf for GFP expression, using a fluorescence stereomicroscope, and raised until adulthood. Potential founders were then crossed with WT zebrafish and the progeny with consistent GFP expression was selected and raised until adulthood, establishing this way the GFP-USE transgenic line. For the USE-RNAs microinjection, one-cell stage AB WT embryos were microinjected with 3nL containing 300ng/ul USE and USEmt RNAs. Embryos were screened and quantified at 24hpf and 5dpf for GFP expression as described before. The fluorescence was calculated using the following formula: GFP Fluorescence = Integrated Density – (Area of selected area X Mean fluorescence of background readings).

For the assessment of the impact of the rs3087967 risk variant, we performed *in vivo* mosaic transgenesis assay. One-cell stage AB WT embryos were microinjected with 3nL containing 25ng/ul Tol2 transposase mRNA and 25ng(ul phenol/chloroform-purified reporter vectors (pUC19-miniTOL-GFP normal variant and pUC19-miniTOL-GFF risk variant and mylz:mCherry). Embryos were raised until reach the 4dpf larval stage, fixed and stained with DAPI (PUREBLU™ DAPI (1351303; Bio-Rad). The images were acquired using SP5II confocal microscope.

### Immunohistochemistry

Transgenic GFP-USE zebrafish embryos were fixed in 4% formaldehyde in PBS1x, overnight at 4°C, and then washed in PBS-T (0.1% Triton X-100 in PBS1x), permeabilized in PBS-T 0.5%, rewashed and incubated with DAPI (1:1000) (Invitrogen) diluted in PBS-T 0.1%, for 4h. Embryos were extensively washed and mounted in microscopic slides in 50% glycerol in PBS1x. Images were acquired using Leica SP5II confocal microscope and analyzed in ImageJ. USE and USEmt RNAs microinjected embryos were fixed and washed as described before. For permeabilization, the embryos were incubated with 100% cold MeOH and blocked with in situ hybridization buffer. The embryos were then incubated with anti-cleaved caspase 3 pAb (1:100) (Anti-caspase-3, cleaved (Ab-2) Rabbit pAB (PC679; Merck) followed by a wash and a second incubation with anti-rabbit 568. Finally, the embryos were washed, the images acquired in Leica M205.and analysed in ImageJ.

pUC19-miniTOL-GFP normal variant and pUC19-miniTOL-GFF risk variant microinjected embryos were fixed, washed and incubated with DAPI as described before. Images were acquired using Leica SP5II confocal microscope and analyzed in IMARIS software (Imaris (RRID:SCR_007370)).

### Cell culture and transfection

HeLa adherent cell line was grown and maintained in Dulbecco’s Modified Eagle Medium (DMEM) with GlutaMAX, supplemented with 10% fetal bovine serum (FBS) and 1% penicillin-streptomycin solution (10000 U/mL) (Gibco, Thermo Fisher Scientific). The cell line was maintained in a humidified incubator at 37° C with 5% CO_2_. HeLa cells were transfected with 500 ng of plasmid DNA using Lipofectamine 2000 (Thermo Fisher Scientific) and following manufacturer’s instructions. To knockdown *PTBP1*, two siRNAs targeting this RBP (siPTBP1) were transfected into HeLa cells (siPTBP1_1: 5’- GCACAGUGUUGAAGAUCAU-3’; siPTBP1_2: 5’- AACUUCCAUCAUUCCAGAGAA-3’; Sigma-Aldrich), using the jetPRIME® (Polyplus Transfection) reagent, following the manufacturer’s guidelines. A non-target siRNA pool (siNTC: 5’- UGGUUUACAUGUCGACUAA-3’; 5’- UGGUUUACAUGUUGUGUGA-3’; 5’- UGGUUUACAUGUUUUCUGA-3’; 5’- UGGUUUACAUGUUUUCCUA-3’; Dharmacon) was also transfected, to serve as a negative control. Cells were transfected at 50% confluency with 50 nM of siPTBP1 (25 nM siPTBP1_1 + 25 nM siPTBP1_2) or 50 nM of siNTC. Cells were collected for total RNA extraction 48h post-transfection.

### RNA extraction and RT-qPCR

RNA was extracted using TRIzol (Ambion), according to the manufacturer protocol. 1 µg of RNA were treated with Recombinant DNase I (Roche) and cDNA was synthesized using Superscript IV reverse transcriptase (Invitrogen) and random hexamers. RT-qPCR reactions were performed using SYBR Select Master Mix (Applied Biosystems) in the 7500 Fast Real-Time PCR System thermocycler (Applied Biosystems). The list of primers used are listed in Supplementary Table 1 and the results were analysed using the ΔΔCt method using 18S as the housekeeping gene.

### Statistical Analysis

Statistical analyses were performed using unpaired Student’s t-test and by using χ^2^ test with Fisher correction. p-values <0.05 were considered as statistically significant. *, p<0.05; **, p<0.01; ***, p<0.001; ****, p<0.0001.

## Discussion

USEs are characterized as U-rich *cis*-auxiliary elements capable of modulating mRNAs 3’-end formation by acting as an extra-platform for the recruitment of *trans*-acting factors, such as RBPs^5,13,17,18,20,26,37,40,45,63–65^. As a result, USEs are considered to be essential to ensure a proper control of gene activity. These elements were initially described in retrovirus, such as the HIV-1, in the selective use of the PAS at the 3’ of the duplicated viral transcripts^21,24,66^. It was proposed that the existence of USEs in HIV-1 revealed the need for a level of regulation that is not normally required by other PAS. However, this concept was challenged when USEs were identified in cellular genes^14,15,19,22,26,29^ and it is now widely accepted that the general arrangement of mRNA 3’-end formation signals in eukaryotes contains an USE associated with a PAS^67^.

In this work we found that a short motif (DplUSE) located in the USE of the *Drosophila melanogaster’s polo* gene^14^ is also present in highly divergent vertebrate species as zebrafish, mouse and human. Importantly, we show that this motif has conserved functions in gene regulation in the fruit fly, zebrafish and humans, being required to sustain high levels of gene activity in distantly related bilaterian species. Further supporting that this motif may participate in an ultra-conserved mechanism, when analysing the molecular machinery that operates at the DplUSE sequence, we found that several RBPs bind specifically to this motif in human cells and that one of these RBPs is the same orthologue as the one found to bind to and to be required for the DplUSE function in the fruit fly^14^. These results support the existence of a very ancestral mechanism common to highly divergent bilaterians that might have important biological roles. Indeed, this seems to be the case, since the DplUSE sequence on average shows higher levels of conservation in vertebrates than their respective 3’UTRs, suggesting a selective pressure to keep this motive or alternative ones with similar functions in the genome. Nevertheless, as only 31 orthologue genes common to zebrafish, mouse and human have the canonical DplUSE, it is likely the existence of alternative motifs. Supporting this hypothesis, previous studies have identified a similar functional motif in the human gene *prothrombin F2*, diverging in only one nucleotide, comparing to the DplUSE motif (TTGT[G/T]TTT)^29^, and when analysing the conservation of the DplUSE motif across vertebrate species, we observed that some nucleotides are more conserved than others.

DplUSE motif seems to be essential in many biological contexts. In *Drosophila* it was demonstrated that its absence dysregulates the cell cycle gene *polo*, causing severe phenotypes in mitosis in neuroblasts and abdominal segments of the adult fly^14^. In the current work we show that using a dominant negative for the molecular players that operate at the DplUSE sequence, causes severe developmental phenotypes in zebrafish embryos, suggesting that DplUSE controlled genes and the molecular players that operate at the DplUSE are important for vertebrate development. Concomitantly, a similar dominant negative strategy applied to human cells lead to downregulation of 9 out of 9 tested genes (2 of them showing a tendency), from the pool of the 31 DplUSE-containing genes that are common to zebrafish, mouse and humans and that are enriched for human diseases such as Nervous system diseases, Congenital abnormalities, and Malignant Neoplasms. EXT2, for instance, is a glycosyltransferase deregulated in the EXT2-related syndrome where one of the manifestations of the disease is microcephaly [73], a phenotype partially recapitulated in zebrafish embryos microinjected with *DplUSE* RNA. *CLDN12* (Claudin12) is another DplUSE-containing gene, which is overexpressed in colorectal carcinomas^68^. Further suggesting that DplUSE controlled genes have important relevant functions to human health, we found that the 2110 human DplUSE containing genes are enriched for several diseases, including Congenital Abnormalities, Colorectal Cancer and Malignant Neoplasm. These results lead us to postulate that alterations in the DplUSE sequence might be at the basis of the development of human disease. This is the case of the colorectal cancer associated SNP rs3087967, located at the 3’UTR of the gene POU2AF2/C11orf53 known to have a tumorigenic potential, and which mechanistic association to colorectal cancer development has not yet been understood. We show that rs3087967 generates an ectopic DplUSE motif that enhances mRNA activity in intestine cells *in vivo*, being a reasonable mechanistic explanation for the involvement of rs3087967 in colorectal cancer development.

Overall, our study has unveiled the emerging importance of the DplUSE motif, identified in *Drosophila*, and that now we show to have a conserved function across vertebrates and that likely represents an ultra-conserved mechanism of gene expression control. We also show that this motif and the associated ultra-conserved mechanism of gene regulation is of extreme importance to understand several human diseases.

## Acknowledgements

We thank all the current and past members of the Gene Regulation and the Vertebrate Development and Regeneration research groups for their contributions and critical discussions. We are very grateful to Gideon Dreyfuss (Howard Hughes Medical Institute, University of Pennsylvania School of Medicine, USA) and Chris Smith (Cambridge University, UK) for kind gifts of antibodies. The laboratory of AMoreira was funded by National Funds through FCT - Fundação para a Ciência e a Tecnologia, I.P., under the project UIDB/04293/2020, by Programa Gilead Génese, ref 13854, by FCT under the project EXPL/SAU-PUB/1073/2021 (http://doi.org/10.54499/EXPL/SAU-PUB/1073/2021) and by Programa Operacional Regional do Norte and co-funded by European Regional Development Fund under the project “The Porto Comprehensive Cancer Center” with the reference NORTE-01–0145-FEDER-072678— Consórcio PORTO.CCC—Porto Comprehensive Cancer Center. The laboratory of JBessa was supported by: European Research Council (ERC) under the European Union’s Horizon 2020 research and innovation programme (ERC-2015-StG1077 680156-ZPR); “La Caixa” Foundation (under the grant agreement HR21-01212); Portuguese funds through Fundação para a Ciência e a Tecnologia (FCT) in the framework of the project PTDC/BIA-MOL/3834/2021. JB is supported by the FCT grant CEECIND/03482/2018. Ana Eufrásio and Joana Azevedo are PhD students (FCT studentships SFRH/BD/147762/2019 to AEufrásio and 2022.14339.BD to JAzevedo). IP-C is supported by a FCT Researcher contract (DL 57/2016/CP1355/CT0016).

## Author contributions

AEufrásio, AMoutinho, JAzevedo and AJesus performed all the zebrafish experiments; JMachado performed all the experiments in human cell lines and IP-C supervised the work; JTavares designed the bioinformatic script, AFerreira, FHenriques, JTeixeira and AEufrasio performed all the bioinformatic analyses; PBPP performed the in vitro transcription and UV crosslinking assays; JBessa, AMoutinho, AEufrásio, PBPP and AMoreira wrote the paper; JBessa and AMoreira designed the project and supervised the team.

---

**Figure S1** - (A) Representative graph showing the percentage of the 31 ortholog DplUSE- containing genes that are associated with the respective disease (DisGenet data source). (B) Representative graph of the extracted conservation values from each nucleotide within the USE and 3’UTR sequence, for the 31 ortholog genes (data source: 100vertebrates) (C) Representative grpah of the extracted conservation values from each nucleotide within the USE and 3’UTR sequence, for the 2110 human genes (data source: 100vertebrates) (D) Representative graph showing the average conservation value for each nucleotide within the 31 DplUSE containing genes. (E) Discovered motifs found from the alignment of the converted DplUSEs from the 31 ortholog genes, in mouse, zebrafish and american alligator. (F) Discovered motifs found from the alignment of the converted DplUSEs from the 2110 human genes, in mouse, zebrafish and american alligator. (G) DplUSE-containing genes GO terms from human, mouse (H) and zebrafish (I). (J) Venn diagram representing the common GOs highly enriched between human, zebrafish and mouse (F.E≥1.5). A total of 10 common GO terms were found between human and mouse, 4 between zebrafish and mouse and 2 between human and zebrafish. (K)Table representing the GO terms found to be highly enriched in human, zebrafish and mouse.

**Figure S2** - Microinjection of DplUSE RNA causes a decrease in GFP expression and microphthalmia in transgenic GFP-DplUSE animals, at 48hpf. (A) Representative image of GFP expression in transgenic GFP-DplUSE zebrafish embryos not microinjected and microinjected with USE RNA or USEmt, at 48hpf. Images were acquired using Leica M205.(B) Quantification of GFP expression in notochord (n=37 DplUSE; n=57 DplUSEmt).(C) Quantification of the percentage of transgenic GFP-USE embryos that showed a patchy expression of GFP in the notochord (68%,n=79 DplUSE; 22%,n=61 DplUSEmt). (D) Quantiffication of the eye size (n=53 DplUSE and n=47 DplUSEmt) and (E) microphthalmia (28%, n=83 DplUSE; 9%, n=91 DplUSEmt) of transgenic GFP-USE embryos upon the microinjection of DplUSE RNA and DplUSEmt RNA, at 48hpf. Statistical significance was determined by χ2 test with Fisher correction or by two tailed unpaired t-test. ***p<0.001; **p<0.01; *p<0.05; ns p>0.05. (Scale bar = 100μm).

**Figure S3** - Microinjection of GFP-DplUSE RNA in one-cell stage embryos leads to the development of various abnormalities 5 days upon the microinjection procedure.(A) Quantification of the several defects developed 5 days upon the microinjection of GFP-DplUSE RNA (in green) and GFP-DplUSEmt RNA (in grey) in one-cell stage zebrafish embryos. Statistical significance was determined using χ2 with Fisher correction. (B) Pie chart representing the total number of defects developed upon the microinjection of GFP-DplUSE RNA and GFP-DplUSEmt RNA. (C) Representative images showing the various abnormalities upon microinjection of GFP-DplUSE RNA at 5 dpf, indicated with red arrows. Scale bar = 100 μm. Statistical significance was determined by χ2 test with Fisher correction or by two-tailed unpaired t-test. ***p<0.001; **p<0.01; *p<0.05; ns p>0.05 Images acquired with Leica M205.Scale bar = 100 μm.

**Figure S4** - To map the binding sites in the DplUSE, several RNAs were in vitro transcribed: A) DplUSEmt-2, DplUSE-5’mt, DplUSE-3’mt, DplUSE-(a), DplUSE-(b) and DplUSE-(c). B-C) UV. cross-linking assays were performed using these RNAs with HeLa cell nuclear extracts. As a positive and negative controls an DplUSE RNA, containing the wild-type DplUSE sequence (lanes 1) and a RNA transcribed from a linearized pGEM7 polylinker (lane 5) were used, respectively.

**Figure S5** - PTBP1 gene expression was efficiently silenced in HeLa cells. In order to knockdown PTBP1, HeLa cells were transfected with two siRNAs targeting PTBP1. A siNTC was also transfected, to act as a negative control. mRNA levels of PTBP1 were assessed by RT-qPCR, showing a mean knockdown of ∼75%.

**Supplementary table 1**. (A) Oligonucleotides and (B) antibodies used in this study.

