## Supplemental figures for "A short 3’UTR motif regulates gene expression in bilaterians"

### Figure S1

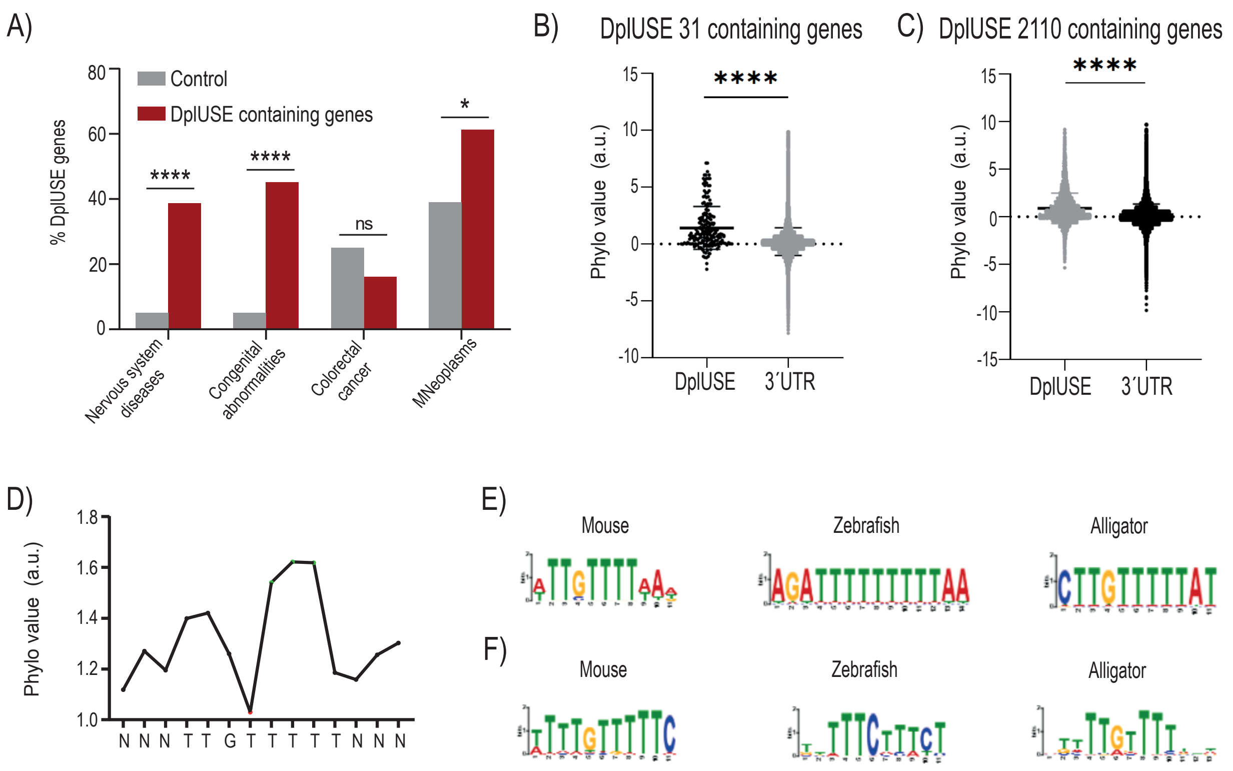


G)

| *Homo sapiens* DpIUSE-containing genes GO Terms | Number of genes | Fold enrichment | FDR |
|---|---|---|---|
| import into nucleous (GO:0051170) | 24 | 2.33 | 4.53E-02 |
| homophilic cell adhesion via plasma membrane adhesion molecules (GO: 0007156) | 35 | 2.14 | 1.65E-02 |
| regulation of mRNA processing (GO:0050684) | 29 | 2.13 | 4.79E-02 |
| regulation of organelle assembly (GO:1902115) | 39 | 2.11 | 1.12E-02 |
| cell-cell adhesion via plasma-membrane adhesion molecules (GO:0098742) | 48 | 1.9 | 1.29E-02 |
| positive regulation of cell migration (GO:0030335) | 89 | 1.74 | 7.50E-04 |
| regulation of Wnt signalling pathway (GO:0030111) | 54 | 1.71 | 4.42E-02 |
| response to growth factor (GO:0070848) | 80 | 1.69 | 5.06E-03 |
| regulation of cell migration (GO:0030334) | 147 | 1.67 | 1.42E-05 |
| positive regulation of cell motility (GO:2000147) | 89 | 1.66 | 3.29E-03 |
| cell-cell adhesion (GO:0098609) | 84 | 1.66 | 4.74E-03 |
| cellular response to growth factor stimulus (GO:0071363) | 74 | 1.66 | 1.24E-02 |
| covalent chromatin modifications (GO:0016569) | 59 | 1.66 | 4.61E-02 |
| mRNA processing (GO:0006397) | 70 | 1.64 | 2.52E-02 |
| positive regulation of locomotion (GO:0040017) | 90 | 1.63 | 5.10E-03 |

H)

| *Mus musculus* DpIUSE-containing genes GO Terms | Number of genes | Fold enrichment | FDR |
|---|---|---|---|
| multivesicular body sorting pathway (GO:0071985) | 11 | 3.79 | 3.59E-02 |
| cellular response to glucose stimulus (GO:0071333) | 17 | 3.75 | 2.28E-03 |
| cellular response to hexose stimulus (GO:0071331) | 17 | 3.62 | 3.11E-03 |
| cellular response to monosaccharide stimulus(GO:0071326) | 17 | 3.55 | 3.65E-03 |
| cellular response to carbohydrate stimulus (GO:0071322) | 18 | 3.29 | 4.74E-03 |
| Golgi to plasma membrane transport (GO:0006893) | 14 | 3.15 | 3.35E-02 |
| cellular glucose homeostasis (GO:0001678) | 21 | 3.11 | 2.74E-03 |
| regulation of epithelial to mesenchymal transition (GO:0010717) | 22 | 2.77 | 6.58E-03 |
| response to glucose (GO:0009749) | 22 | 2.63 | 1.50E-02 |
| post-Golgi vesicle-mediated transport (GO:0006892) | 20 | 2.6 | 2.97E-02 |
| response to hexose (GO:0009746) | 22 | 2.55 | 1.87E-02 |
| response to monosaccharide (GO:0034284) | 22 | 2.52 | 2.03E-02 |
| response to carbohydrate (GO:0009743) | 23 | 2.32 | 4.03E-02 |
| vesicle-mediated transport to the plasma membrane (GO:0098876) | 23 | 2.26 | 4.89E-02 |
| establishment of vesicle localization (GO:0051650) | 27 | 2.23 | 2.84E-02 |

I)

| *Danio rerio* DpIUSE-containing genes GO Terms | Number of genes | Fold enrichment | FDR |
|---|---|---|---|
| positive regulation of protein ubiquitination (GO:0031398) | 10 | 6.74 | 2.80E-03 |
| ribonucleoside metabolic process (GO:0009119) | 7 | 6.44 | 3.63E-02 |
| positive regulation of protein modification by small protein conjugation or removal (GO:1903322) | 10 | 6.13 | 4.20E-03 |
| regulation of protein ubiquitination (GO:0031396) | 12 | 5.17 | 3.28E-03 |
| regulation of protein modification by small protein conugation or removal (GO:1903320) | 13 | 5.06 | 2.27E-03 |
| cytosolic transport (GO:0016482) | 12 | 3.62 | 3.62E-02 |
| Notch signalling pathway (GO:0007219) | 13 | 3.25 | 4.72E-02 |
| endocrine system development (GO:0035270) | 18 | 3.09 | 1.10E-02 |
| connective tissue development (GO:0061448) | 20 | 2.83 | 1.29E-02 |
| cartilage development (GO:0051216) | 19 | 2.83 | 1.79E-02 |
| protein polyubiquitination (GO:0000209) | 20 | 2.56 | 4.76E-02 |
| organelle fission (GO:0048285) | 22 | 2.5 | 3.13E-02 |
| small molecule biosynthetic process (GO:0044283) | 34 | 2.21 | 8.38E-02 |
| mitotic cell cycle process (GO:1903047) | 30 | 2.17 | 2.60E-02 |
| nucleobase-containing compound biosynthetic process (GO:0034654) | 49 | 2.13 | 1.37E-03 |

J)

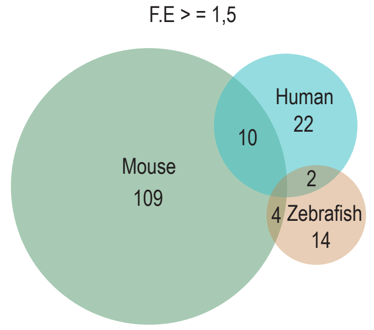


K)

| | Human | Mouse | Zebrafish |
|---|---|---|---|
| positive regulation of cell migration (GO:0030335) | • | • | |
| mRNA processing (GO:0006397) | • | • | |
| positive regulation of cell motility (GO:2000147) | • | • | |
| response to growth factor (GO:0070848) | • | • | |
| positive regulation of locomotion (GO:0040017) | • | • | |
| cell division (GO:0051301) | • | • | |
| positive regulation of cellular component movement(GO:0051272) | • | • | |
| cellular response to growth factor stimulus (GO:0071363) | • | • | |
| chromatin organization (GO:0006325) | • | • | |
| mRNA metabolic process (GO:0016071) | • | • | |
| chromosome organization (GO:0051276) | | • | • |
| establishment of localization in cell (GO:0051649) | | • | • |
| cellular protein localization (GO:0034613) | | • | • |
| cellular macromolecule localization (GO:0070727) | | • | • |
| mitotic cell cycle (GO:0000278) | • | | • |
| mitotic cell cycle process (GO:1903047) | • | | • |

### Figure S2

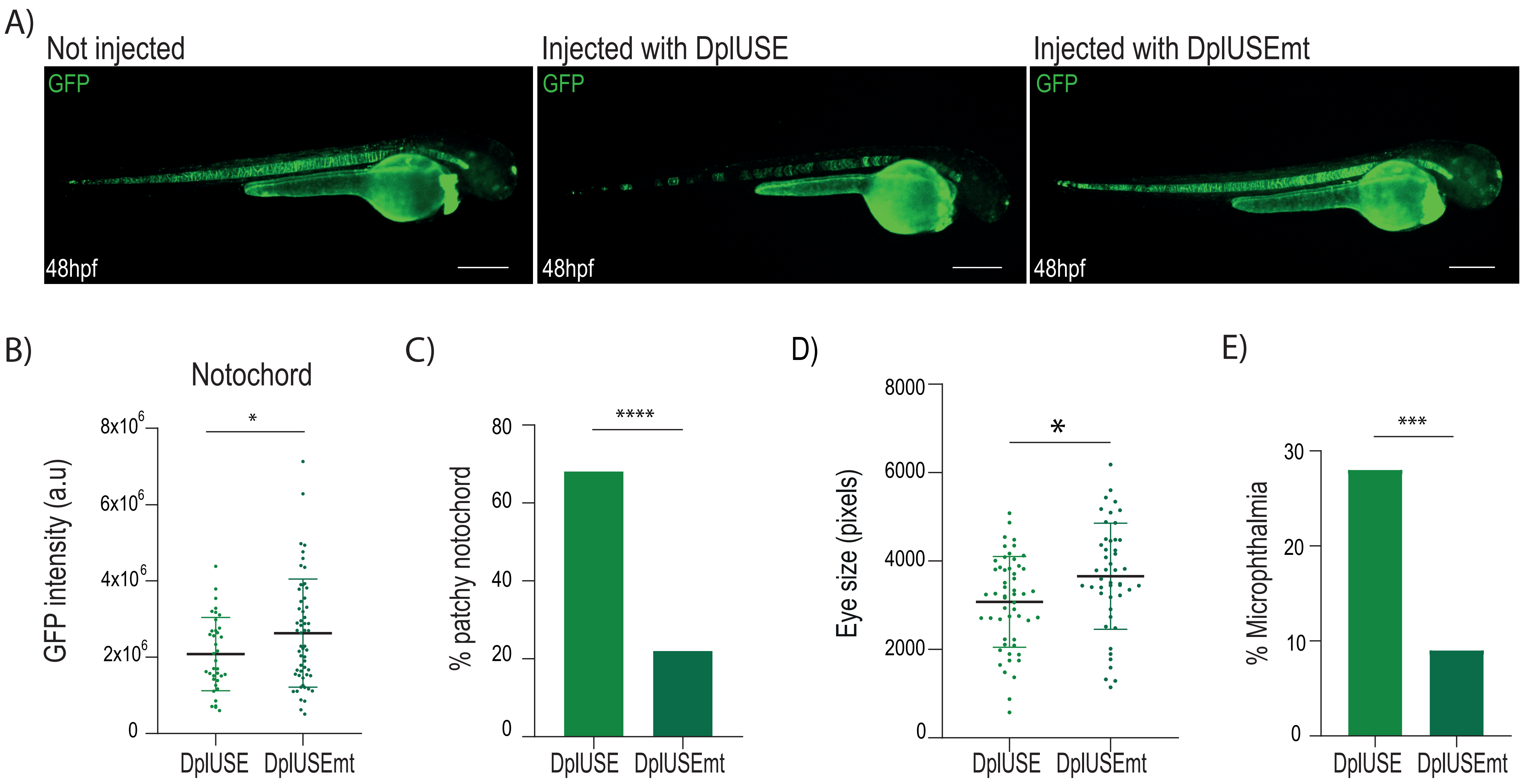
A)
Not injected
Injected with DplUSE
Injected with DplUSEmt
GFP
48hpf
B)
Notochord
GFP intensity (a.u)
8x10^6
6x10^6
4x10^6
2x10^6
0
*
DplUSE
DplUSEmt
C)
% patchy notochord
80
60
40
20
0
****
DplUSE
DplUSEmt
D)
Eye size (pixels)
8000
6000
4000
2000
0
*
DplUSE
DplUSEmt
E)
% Microphthalmia
30
20
10
0
***
DplUSE
DplUSEmt

Figure S3

A) Defects on 5dpf zebrafish embryos

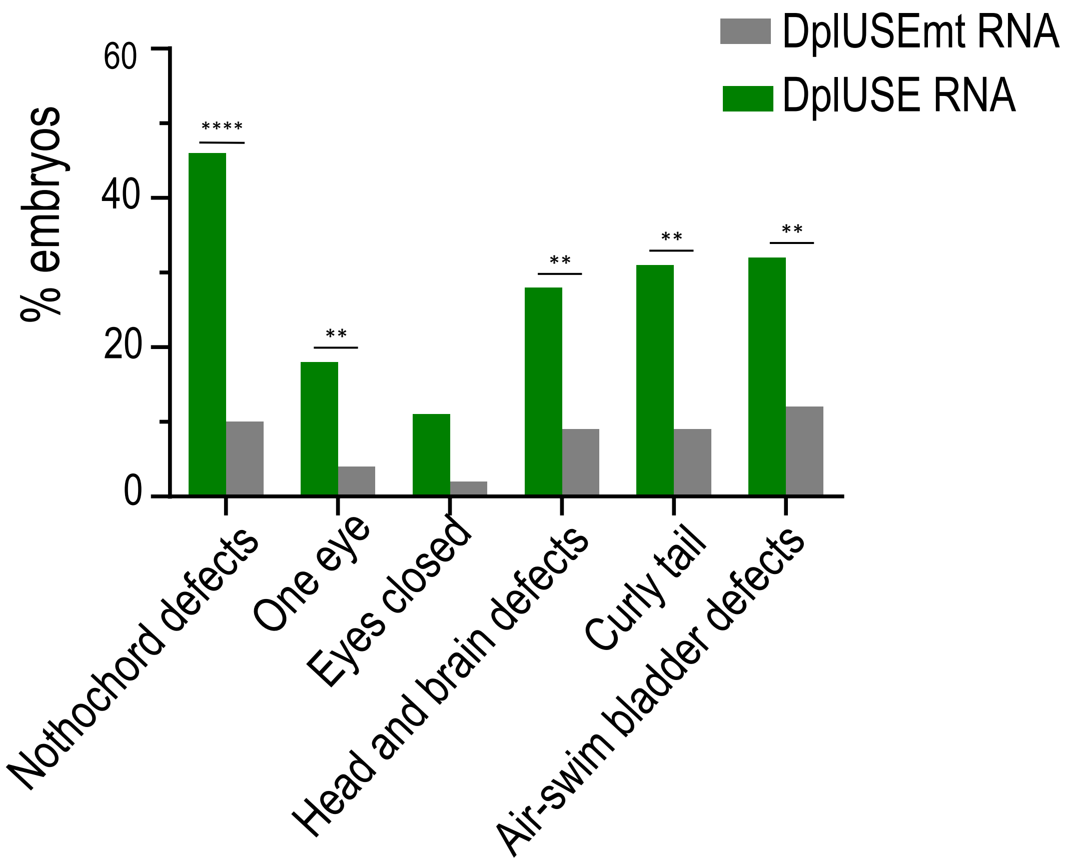


B) Total number of defects in 5dpf zebrafish embryos

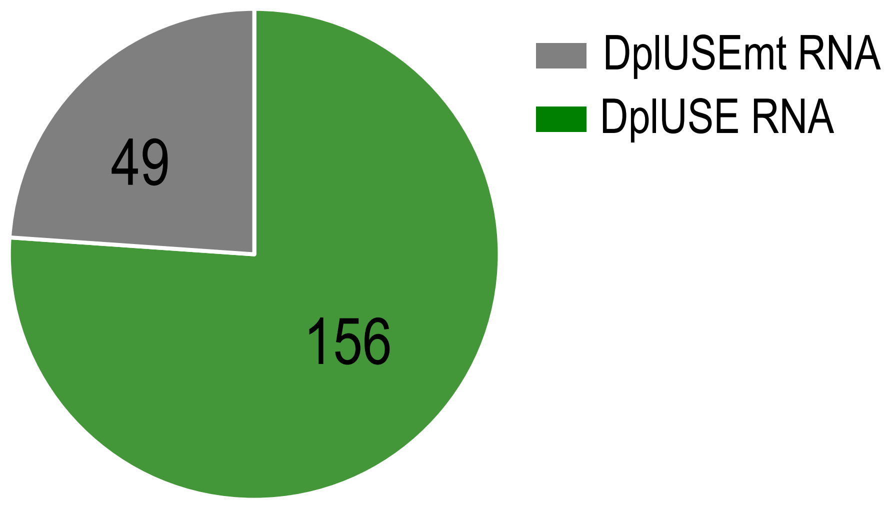


C)

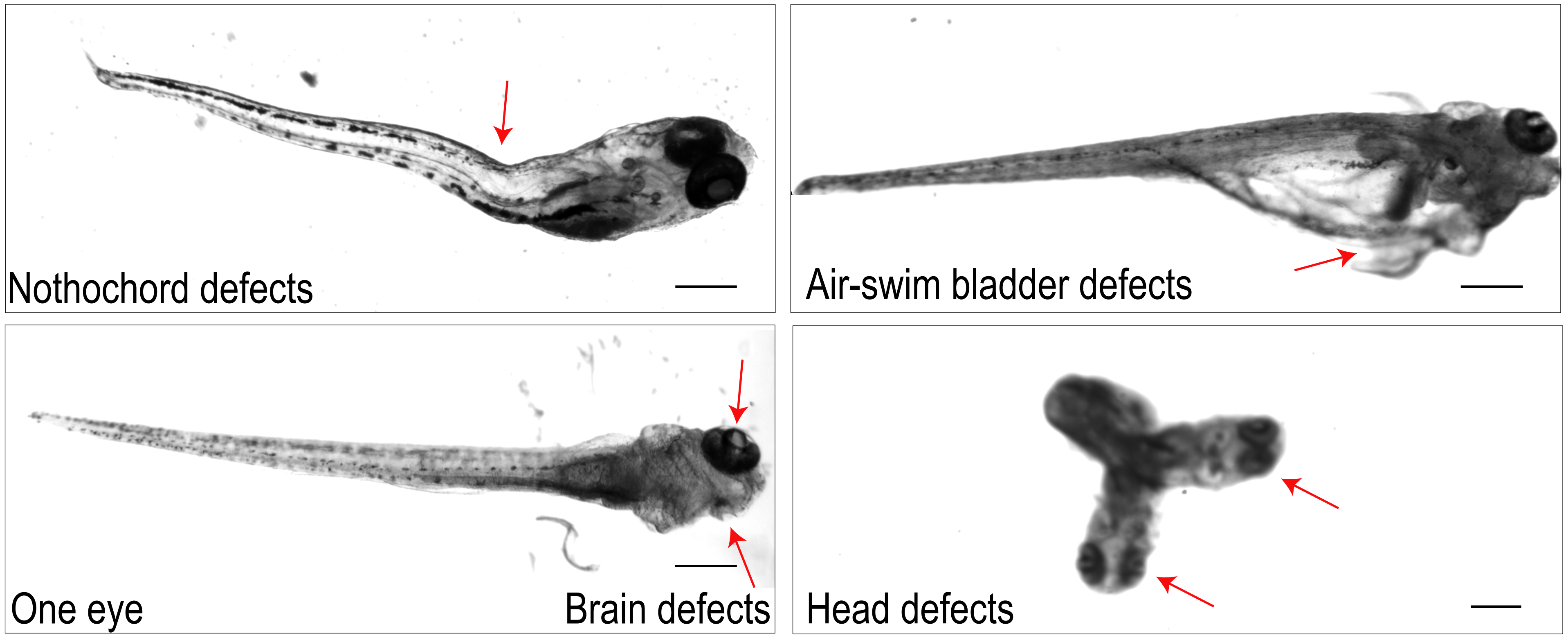

Figure S4

A)

| | |
|---|---|
| **DplUSE** | AAUUUAUUUGUUUUUGCCCCUUCCCCUU |
| **DplUSEmt-2** | AAUUCAUAGACGACAAACGCAAGAUCUU |
| **DplUSE-5`mt** | AAUUCAUAGACGACAGCCCCUUCCCCUU |
| **DplUSE-3`mt** | AAUUUAUUUGUUUUUAACGCAAGAUCUU |
| **DplUSE-(a)** | AAUUUAUUUGUUUUU |
| **DplUSE-(b)** | GUUUUUGCCCC |
| **DplUSE-(c)** | GCCCCUUCCCCUU |

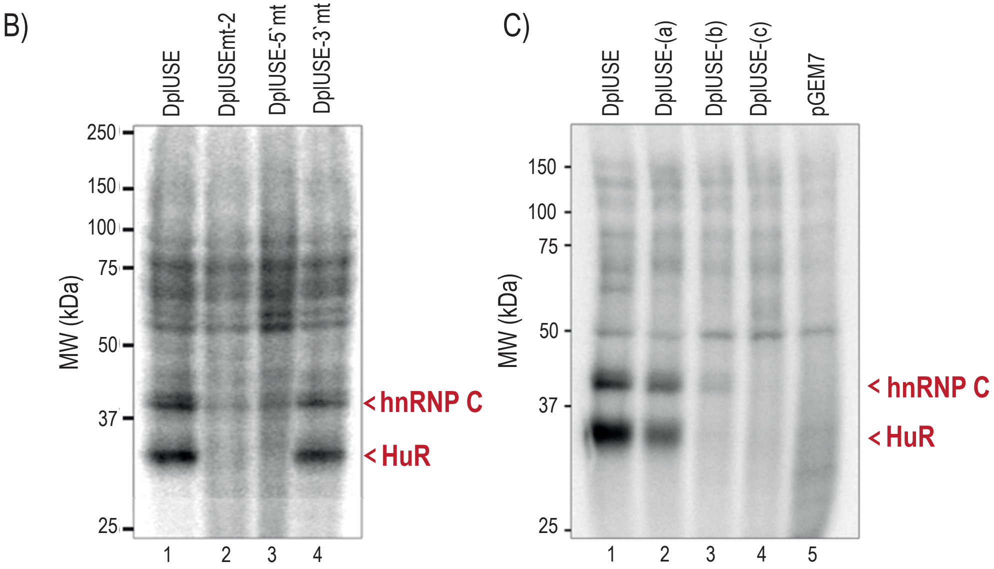

Figure S5

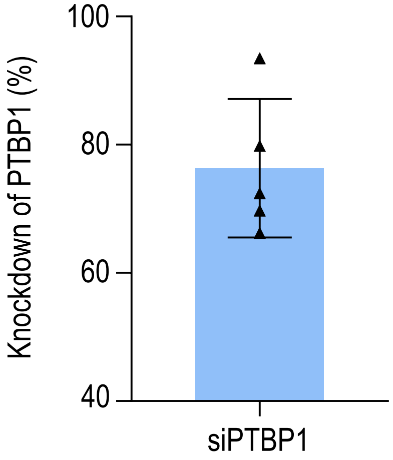

A)

| Oligonucleotides | Sequence |
|---|---|
| FW_useORF | CCGCTCGAGTAATAAGGCCAAATGACTTCTGGG |
| RV_useORF | CGGGGTACCACAAAGAGCAGCATTGAGG |
| FW_useORF_SiteDirectMut | CCTAAAACAATCTTTGGAATACACC |
| RV_useORF_SiteDirectMut | GGTGTATTCCAAAGATCGTTTTAGG |
| CSNK1A1_F | AAAGCAGAAAGCAGCACAGCAG |
| CSNK1A1_R | TTGTCAGTTTGCTTGCCTGTGG |
| KIF20A_F | GCAATGAACGGGGAATTGGC |
| KIF20A_R | GGATGCCTGTCCCACTTCTG |
| MAP4K5_F | CAGACAGGGTTGTCGTTTTGG |
| MAP4K5_R | CATGTCCAGCCAAGATGTAGAGA |
| CLDN12_F | TGGCAACCATTGTACTCCCA |
| CLDN12_R | TTCAATGGCAGAGAGGCGAG |
| EXT2_F | GACCTCTGACGAGCTGCAAT |
| EXT2_R | GTCCCAGAGATGCAGACGAC |
| MED18_F | CAAGCCAGCCCATTTGTTCTC |
| MED18_R | TGTCTCCCATTTCTGGCTGTC |
| DKC1_F | TATGTGGGGATTGTCCGGCT |
| DKC1_R | CTGCAGCAATAAGTGGGGGT |
| MBNL2_F | GGCCGTTGTTCGAGAGAGAA |
| MBNL2_R | AAGCATTGCTGCTGCAGTTT |
| NR5A2_F | CCCTATACCAGCTCACCCGA |
| NR5A2_R | AGATGTGGGATGCTTGCTGG |
| eGFP_F | GAGCTGAAGGGCATCGACTT |
| eGFP_R | TTCTGCTTGTCGGCCATGAT |
| PTBP1_F | TCCAGAAGGACCGCAAGATG |
| PTBP1_R | TCGTGGTTGTGCAGGTCAAT |

B)

| Antibodies |
|---|
| Anti-caspase-3, cleaved (Ab-2) Rabbit pAB (PC679 - Merck) |
| PUREBLU™DAPI (1351303 - Bio-Rad) |

**Supplementary Table 1. (A)** Oligonucleotides and **(B)** antibodies used in this study
